# A mechanistic classification of *GNAO1* mutations links Gβγ sequestration to hyperkinetic disease and treatment response

**DOI:** 10.64898/2026.10.24.761540

**Authors:** Gabrielle M. Soden, Amy Robichaux-Viehoever, Kaitlin M. Soden, Jeff P. Laux, Henrik G. Dohlman, Daniel J. Christoffel

## Abstract

Gα_o_ is the most prevalent G protein α subunit in the brain and is critical for signal transduction through G protein-coupled receptors. Mutations in the gene encoding Gα_o_ (*GNAO1*) cause a rare neurodevelopmental disorder with involuntary movements (NEDIM) and, in some patients, developmental and epileptic encephalopathy (DEE). Here, we integrate human genetic analysis with cellular assays to define mechanistic classes of *GNAO1* mutations and *in vivo* approaches to evaluate therapeutic strategies. We grouped variants into two functional categories: those predicted to sequester Gβγ and those with reduced Gα_o_ expression and therefore increased Gβγ signaling. Analysis of patient data revealed that mutations predicted to impair Gβγ release are significantly associated with NEDIM. Using the *GNAO1*^R209H/+^ mouse as a model of NEDIM, we evaluated two small molecules: caffeine, a nonselective adenosine receptor antagonist, and gallein, a selective modulator of Gβγ–effector interactions. Both compounds significantly reduced hyperlocomotion in the open field test. Whole-cell patch-clamp electrophysiology revealed sex- and cell type–specific hyperexcitability in spiny projection neurons within the dorsolateral striatum of *GNAO1*^R209H/+^ mice. Together, these findings implicate dysfunction of dorsolateral striatal pathways in GNAO1-related disorder and suggest that mechanistic understanding of disease variants can guide targeted therapeutic strategies.

## Introduction

G protein–coupled receptors (GPCRs) are critical mediators of cellular signaling, transducing extracellular stimuli such as light, hormones, and neurotransmitters into intracellular signaling cascades. Because of their central role in physiology, GPCRs are targets of widely used drugs, including the adenosine receptor antagonist caffeine. Upon GPCR activation, the heterotrimeric G protein complex, composed of Gα, Gβ, and Gγ subunits, is recruited to the receptor. Activated receptors promote the exchange of GDP for GTP on the Gα subunit, causing dissociation of Gα from the Gβγ dimer. Both Gα and Gβγ can then interact with downstream effectors to initiate intracellular signaling pathways. Signal termination occurs when the intrinsic GTPase activity of Gα converts GTP to GDP, allowing reassembly of the heterotrimer. Disruptions at any step in this signaling cycle can affect cellular communication, and mutations in G protein subunits have been linked to multiple disease states, including neurodevelopmental disorders and cancer.^1–4^

GNAO1-related disorder is a rare neurodevelopmental condition caused by mutations in the gene encoding the Gα_o_ protein (*GNAO1*).^1^ Gα_o_ is the most abundant G protein α subunit in the brain^5^ and plays an important role in neuronal signaling via its binding partner, Gβγ which can modulate various ion channels.^6,7^ Pathogenic variants in *GNAO1* are associated with two major clinical presentations: neurodevelopmental disorder with involuntary movements (NEDIM; OMIM #617493) and the more severe developmental and epileptic encephalopathy (DEE; OMIM #615473), characterized by movement abnormalities and refractory epilepsy.^1,8^

Despite the increasing number of identified *GNAO1* variants, genotype–phenotype relationships remain poorly defined for most mutations.^8^ One notable mutational hotspot occurs at arginine 209 (R209). Structural modeling and cellular assays indicate that substitutions at this residue impair Gβγ release, thereby blunting downstream signaling.^9^ A heterozygous mouse model harboring the R209H mutation (*GNAO1*^R209H/+^) recapitulates key features of the NEDIM phenotype, including pronounced hyperactivity without seizures,^10^ and therefore provides a platform for mechanistic investigation and preclinical therapeutic testing.

Movement disorders often arise from dysfunction within the basal ganglia, including the striatum, a central hub for motor control.^11–13^ Approximately 95% of striatal neurons are spiny projection neurons (SPNs), which are divided into two major populations. Direct pathway SPNs express dopamine D1 receptors coupled to Gα_s/olf_ and can facilitate movement, whereas indirect pathway SPNs express dopamine D2 receptors coupled to Gα_i/o_ and can suppress movement.^14^ Dysregulation of this circuitry is thought to underlie hyperkinesis, including in GNAO1-related disorder.^15–17^ Dopamine D2 receptor antagonists, including haloperidol, and benzodiazepines are commonly used to suppress hyperkinesis but can cause severe adverse effects such as catatonia, limiting their long-term therapeutic utility.^10,18–21^

An alternative therapeutic strategy involves targeting adenosine signaling. Adenosine receptors are co-expressed with dopamine receptors in striatal circuits: D1 SPNs co-express the Gα_i/o_-coupled adenosine A1 receptor, while D2 SPNs co-express the Gα_s/olf_-coupled adenosine A2A receptor.^22^ Adenosine signaling in the striatum is dynamically recruited during movement. Locomotion increases extracellular adenosine and PKA activity in both direct- and indirect-pathway SPNs. Inhibition of striatal PKA signaling in either SPN population impairs locomotor performance, suggesting that normal movement depends on coordinated recruitment of both pathways rather than strictly opponent activation.^23^ These dopamine and adenosine receptor pairs exert opposing control of striatal SPN activity. Furthermore, the nonselective adenosine receptor antagonist caffeine is being explored as a therapeutic in clinical trials for GNAO1 disorder based on experiments in the nematode, *C. elegans*.^24^ Another candidate for therapeutic intervention is gallein, an experimental small molecule that can inhibit interactions between Gβγ and certain downstream effectors.^25^ Gallein is of particular interest because the primary physiological role of Gα_o_ is to tightly regulate Gβγ signaling dynamics.^26^

In this study, we integrate human genetic data, cellular signaling assays, as well as *ex vivo* and *in vivo* approaches to investigate the mechanisms underlying GNAO1-related movement disorders. Building on mechanistic classifications developed by Knight et. al.,^9^ we grouped mutations into two functional categories: those that sequester Gβγ and those that exhibit reduced expression making them unavailable to bind Gβγ leading to increased Gβγ signaling. We find that mutations that sequester Gβγ are much more likely to be NEDIM, while those that create haploinsufficiency are much more likely to be DEE. Using the *GNAO1*^R209H/+^ mouse as a representative NEDIM model, we evaluate the therapeutic potential of caffeine and gallein to reduce hyperlocomotion. Finally, through electrophysiological analysis of striatal neurons, we identify cell-type specific alterations within the dorsolateral striatum, implicating this brain region as a key locus of dysfunction in GNAO1-related disorder. Together, these findings link G protein genotype to phenotype and provide a framework for mechanism-guided therapeutic development.

## Results

### Mechanistic classification of *GNAO1* variants predicts clinical phenotype

GNAO1-related disorder comprises two predominant clinical presentations, DEE and NEDIM. However, the mechanistic basis for these distinct phenotypes remains unclear. We therefore asked whether *GNAO1* mutations could be grouped into functional classes that predict clinical presentation. Building on prior biochemical and cell-based studies, we organized pathogenic variants into two broad categories according to their putative effects on Gβγ availability. This framework predicts that Gβγ-sequestering variants reduce downstream effector signaling and produce a NEDIM phenotype, whereas haploinsufficient variants increase free Gβγ and are associated with DEE (Fig. 1A).

**Figure 1.**
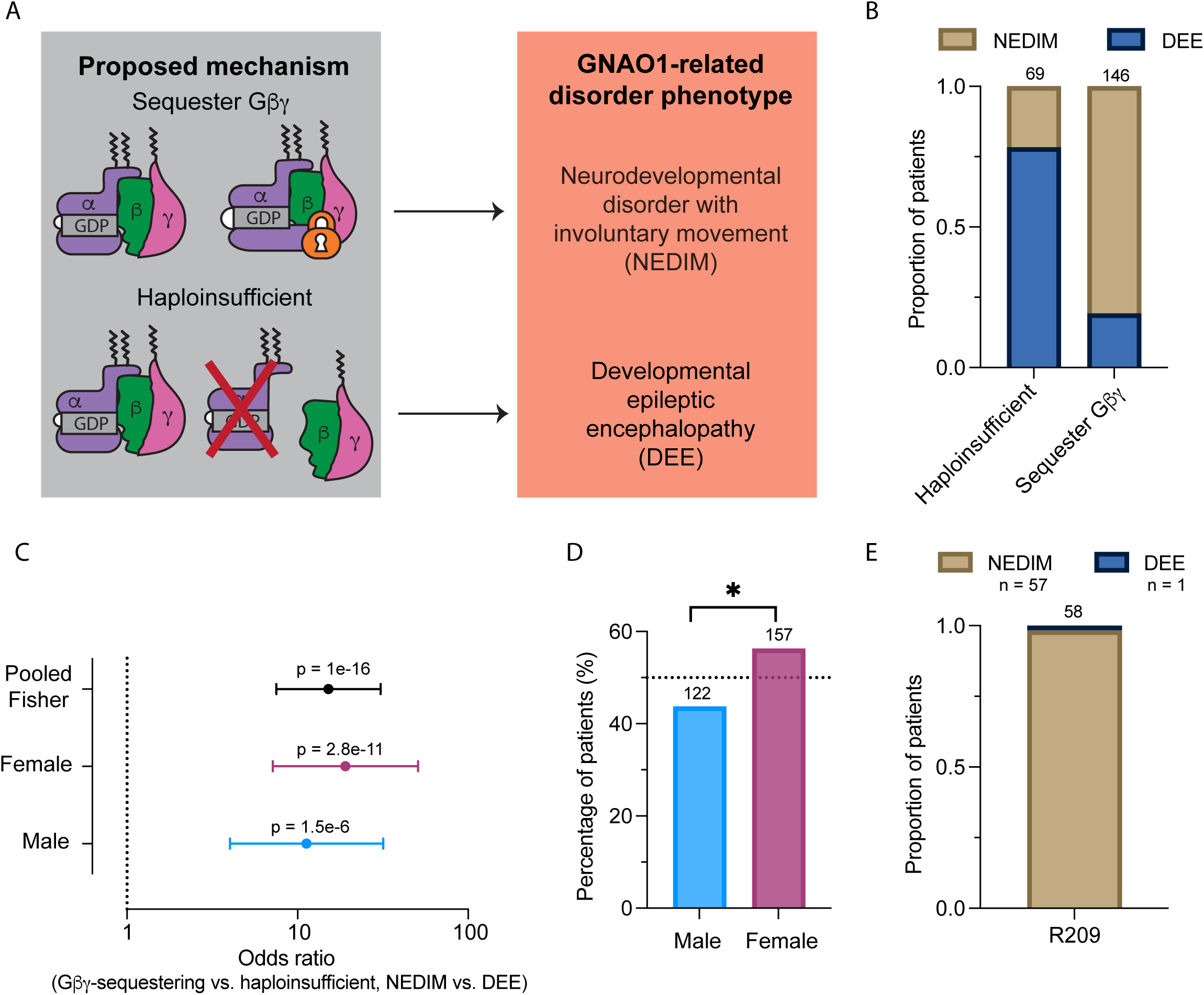
Mechanistic classification of *GNAO1* variants predicts clinical phenotype. (A) Schematic illustrating two proposed mechanistic classes of *GNAO1* pathogenic variants and their predicted clinical outcomes. Gβγ-sequestering variants (top) retain Gα_o_ expression but lock Gα_o_ in an inactive complex, reducing free Gβγ available for downstream signaling, and are predicted to associate with NEDIM. Haploinsufficient variants (bottom) reduce Gα_o_ expression or signaling capacity, increasing relative Gβγ availability, and are predicted to associate with DEE. (B) Proportion of patients classified as DEE (dark blue) or NEDIM (tan) within each mechanistic class. Numbers above the bars indicate the total number of patients per group. (C) Forest plot showing odds ratios (log scale) for NEDIM versus DEE in Gβγ-sequestering compared with haploinsufficient variants, for the pooled cohort and for female- and male-stratified analyses. Points represent odds ratio estimates; horizontal lines represent 95% confidence intervals. Dashed vertical line indicates OR = 1. P-values from Fisher’s exact test. (D) Percentage of GNAO1 disorder patients who are male (n = 122) or female (n = 157) out of 279 total analyzed. Dotted line indicates the expected percentage under a 50:50 sex distribution. Asterisk indicates statistical significance (χ² = 4.39, df = 1, p = 0.036). (E) Proportion of patients carrying any R209 substitution classified as DEE (dark blue, n = 1) or NEDIM (tan, n = 57).

To test this model, we curated published GNAO1-related disorder cases (Table S1), categorized patients as DEE or NEDIM based on clinician-reported phenotypes, and classified variants using criteria adapted from prior BRET and expression studies (Table S2). We used an odds ratio (OR) to assess whether variant mechanism and patient phenotype were related. We found that patients with haploinsufficient variants were significantly more likely to present with DEE whereas patients with Gβγ-sequestering variants were predominantly classified as NEDIM (Fig. 1B). Patients with Gβγ-sequestering variants had 15 times the odds of presenting with NEDIM compared to haploinsufficient variants (95% confidence interval (CI): 7.496 -30.704, Fig. 1C). This association was consistent across sexes: stratified analyses revealed significant enrichment in both females (OR = 19, 95% CI: 7.150 - 50.823 Fig. 1C) and males (OR = 11.2, 95% CI: 4.010 - 31.738, Fig. 1C), and the stratified odds ratios did not differ significantly from one another, arguing against an interaction between sex and mechanism. Logistic regression further confirmed that the mutation mechanism is a strong predictor of phenotype, whereas sex and the sex-by-mechanism interaction terms were not significant (p = .604, p = .471, respectively; Table S3). Together, these results suggest that the molecular mechanism is a major determinant of clinical presentation in GNAO1-related disorder.

Many neurodevelopmental disorders occur predominantly in males.^27–29^ Since we did not have sufficient evidence to establish that sex modified the relationship between the mutation mechanism and the phenotype, we next asked whether sex might instead influence disease prevalence. We compared the observed sex distribution of our 279-patient cohort against a null expectation of a 50:50 male-to-female ratio using a chi-square goodness-of-fit test. Contrary to the pattern seen in other neurodevelopmental disorders, females were significantly overrepresented: 157 patients (56.3%) were female and 122 (43.7%) were male (χ² = 4.39, df = 1, p = 0.036, Fig. 1D). This difference, combined with insufficient evidence for a sex-by-mechanism interaction, further suggests that mutation mechanism is the primary driver of clinical presentation. This aligns with previous work that found a high percentage of GNAO1-related disorder patients were female: an early study reported 12/12 patients were female, and a review of 213 patients reported 56.8% female.^30,31^ These data confirm that the female predominance is not limited to our analysis.

To further evaluate this mechanistic framework, we next focused on the arginine 209 (R209) residue, which is one of the most clinically recurrent mutation sites in GNAO1-related disorder. Among 58 patients carrying R209 substitutions, 57 were classified as NEDIM and only one as DEE, confirming that mutations at this residue are overwhelmingly associated with NEDIM (Fig. 1E). Within this subgroup, R209C was the most common substitution (50%), followed by R209H (36.2%), with R209P (8.6%), R209L (3.4%), and R209G (1.7%) occurring less frequently. These data identify R209 as a mutational hotspot strongly associated with NEDIM and support the use of R209 mutants as a clinically relevant model for investigating GNAO1-related hyperkinetic movement disorder.

### *GNAO1* R209 variants impair G-protein subunit dissociation

Previous work revealed that GNAO1-related disorder mutants fall into two broad mechanistic categories, one where Gα_o_ sequesters Gβγ, and another where Gα_o_ is poorly expressed and fails to bind to Gβγ entirely.^9^ Central to that analysis was the TRUPATH system, a proximity-based bioluminescence resonance energy transfer (BRET) assay in which Gα_o_ is fused to Renilla luciferase (Rluc) and Gγ to green fluorescent protein (GFP, Fig. 2A).^9,32^ This system reports heterotrimer dissociation following agonist addition as a decrease in the GFP/Rluc ratio. Because patients with GNAO1-related disorder present with a movement disorder, we asked whether R209 mutations impair G*α*_o_ coupling at receptors expressed in the brain region most relevant to this phenotype. Accordingly, we used the adenosine A1 (A1R) and dopamine D2 (D2R) receptors, which are highly expressed in the human striatum (https://proteinatlas.org).^33^ We tested R209C and R209H alongside K46E, which is expressed at WT levels but does not respond to agonist and G40R and K278del, which are positive controls for poorly expressed Gα_o_ variants (Fig. S1). When stimulated with their endogenous ligands, the R209C and R209H responses were similar to WT at the A1 receptor but blunted at the D2 receptor. Whereas K278del, K46E, and G40R showed negligible responses at either receptor (Fig. 2B and C, leftmost panel and Fig. S2A, B).

**Figure 2.**
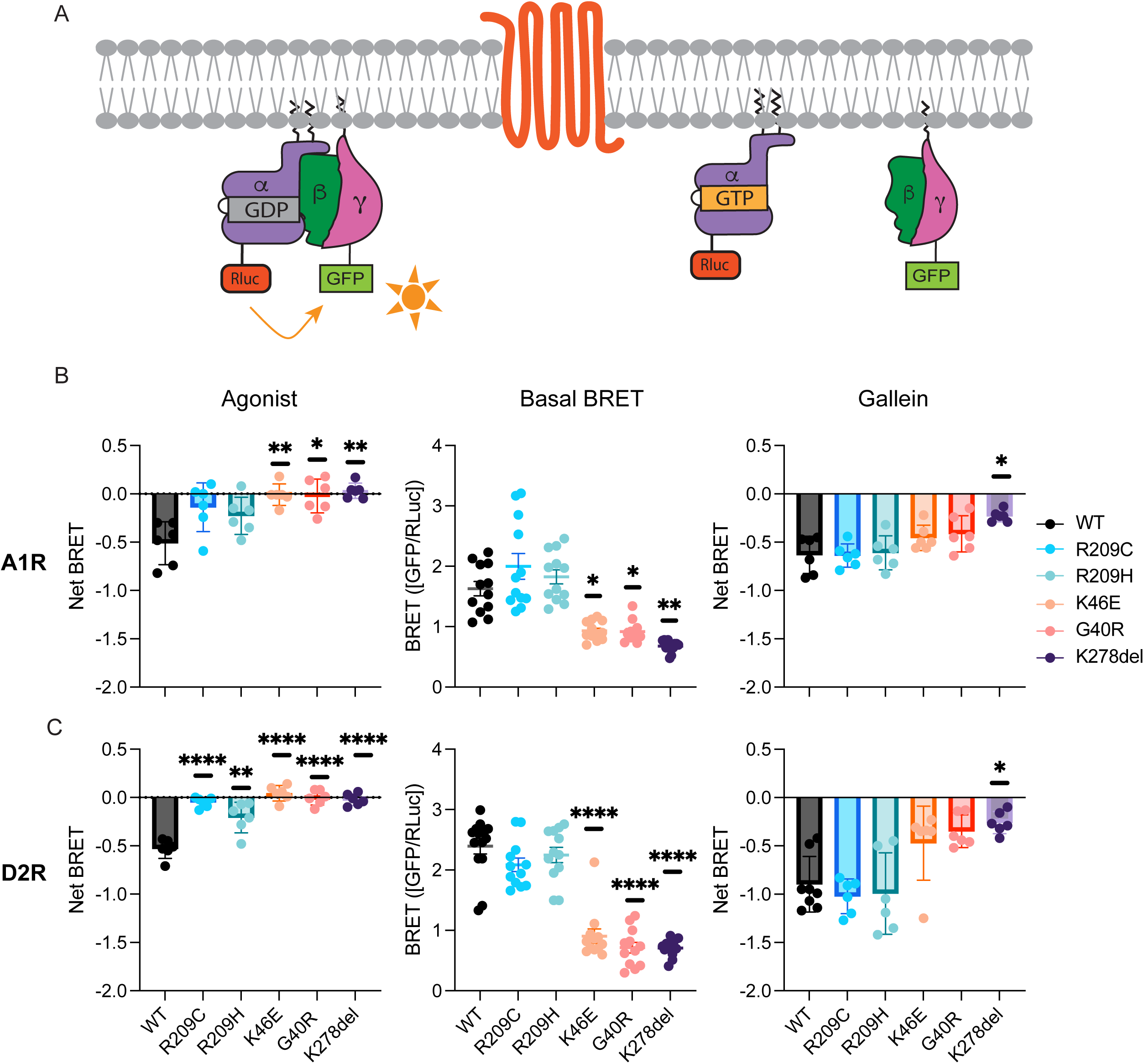
*GNAO1* R209 variants impair Gα_o_ coupling in a receptor-specific manner that is rescued by gallein. (A) Schematic of the TRUPATH BRET assay. In the inactive heterotrimer state, Gα_o_-Rluc and Gβ_3_/Gγ_8_-GFP are in close proximity, producing a high GFP/Rluc BRET signal. Agonist-induced activation causes heterotrimer dissociation, separating Rluc from GFP and reducing the BRET signal. (B–C) Net BRET in response to agonist (left), basal BRET in the absence of agonist (middle), and net BRET in response to the Gβγ inhibitor gallein (right) for the indicated Gα_o_ variants paired with the adenosine A1 receptor (A1R; B) or dopamine D2 receptor (D2R; C). K46E and G40R are non-functional benchmark controls; K278del, a full-codon deletion, is a positive control for poor Gα_o_ expression (Fig. S1). For basal BRET, replicates from agonist and gallein conditions were pooled. Bars represent mean ± SD with individual replicates shown. All experiments have at least 3 biological replicates with 2 technical replicates. Statistical analyses were performed on the mean of the technical replicates from each biological replicate. Statistical comparisons are to WT using one-way ANOVA with Dunnett’s post hoc test, *p < 0.05, ***p < 0.001, ****p < 0.0001.

Basal BRET, measured in the absence of agonist, reflects steady state heterotrimer assembly at the plasma membrane, with lower values indicating reduced Gα_o_ expression or increased constitutive activity. While the agonist data revealed altered subunit dissociation in R209C and R209H, it is not known whether these mutations confer constitutive signaling. We measured basal BRET for each variant at A1R and D2R, pooling replicates from the agonist and gallein conditions (see below), since both were recorded without drug application. Compared with WT, the K278del, K46E, and G40R variants showed markedly reduced basal BRET at both receptors (Fig. 2B and C, middle panel). The reduced basal BRET of K278del and G40R is consistent with poor expression, as confirmed by Western blotting (Fig. S1), whereas K46E has normal expression (Fig. S1), but we hypothesize that the protein may not be trafficked correctly to the membrane. In contrast, R209C and R209H showed basal BRET values comparable to WT at A1R and D2R (Fig. 2B and C, middle panel). Together, these data indicate that R209 mutations cause a receptor-dependent defect in agonist-driven coupling rather than a general failure of heterotrimer formation.

Most drugs used to treat neurological diseases target GPCRs. Our data suggest that the R209H variant disrupts GPCR-driven G-protein signaling in a receptor-dependent manner and, consequently, that pharmacological rescue at the level of the G protein may be appropriate. To that end, we considered the efficacy of gallein, a small molecule that binds Gβγ, as a receptor-independent approach to restoring G protein signaling.^25^ We applied gallein, in the absence of an agonist, to cells expressing the R209C or R209H Gα_o_ variants and either the A1R or D2R receptor. The poorly expressed variants K278del and G40R, as well as K46E, which do not respond to agonist stimulation, were included to assess whether rescue is possible when free Gβγ levels are high. Gallein completely rescued R209C, R209H, K46E, and G40R at A1R and D2R (Fig. 2B and C, rightmost panel and Fig. S2C and D). Only K278del showed partial rescue (Fig. 2B and C, rightmost panel and Fig. S2C, D). We infer that gallein rescues the blunted phenotype by freeing Gβγ from mutant Gα_o_, enabling downstream signaling independently of receptor activation. Given that a single molecule targeting Gβγ can fully rescue two Gα_o_ variants across two receptor systems, our findings position Gβγ as a receptor-independent therapeutic target in GNAO1-related disorder.

### *GNAO1* R209 variants impair Gβγ-effector coupling

The TRUPATH BRET assay (Fig. 2A) reports heterotrimer dissociation through a Gα_o_ subunit fused to Rluc. Because that tag could alter Gα_o_ interactions with its effectors or with its binding partner Gβγ, we used an independent readout that monitors the Gβγ subunit of the heterotrimer. Specifically, we used a complementary BRET assay in which a Gβγ effector fragment (from G protein–coupled receptor kinase, GRK) is fused to the donor NanoLuc (Nluc), and Gβγ is tagged with the acceptor, Venus (Fig. 3A).^34^ Receptor activation induces the release of Gβγ, which can then bind to GRK to produce the BRET signal. Under basal conditions, a well-expressed, functional Gα_o_ should sequester Gβγ, whereas poorly expressed variants should leave Gβγ free to bind GRK. We measured A1R and D2R receptor agonist-induced Gβγ–GRK association, comparing an empty-vector control (EV), WT, R209C, R209H, K46E, G40R, and K278del. At both receptors, R209H and K278del produced an agonist response comparable to WT, whereas EV, R209C, K46E, and G40R showed negligible responses (Fig. 3B and C, leftmost panel). These data show that different R209 substitutions perturb different readouts of the signaling cycle despite converging on the same clinical phenotype. This independent, Gβγ-based assay reproduces the R209C coupling defect and confirms R209H retains coupling, validating the Fig. 2 results with untagged Gα_o_.

**Figure 3.**
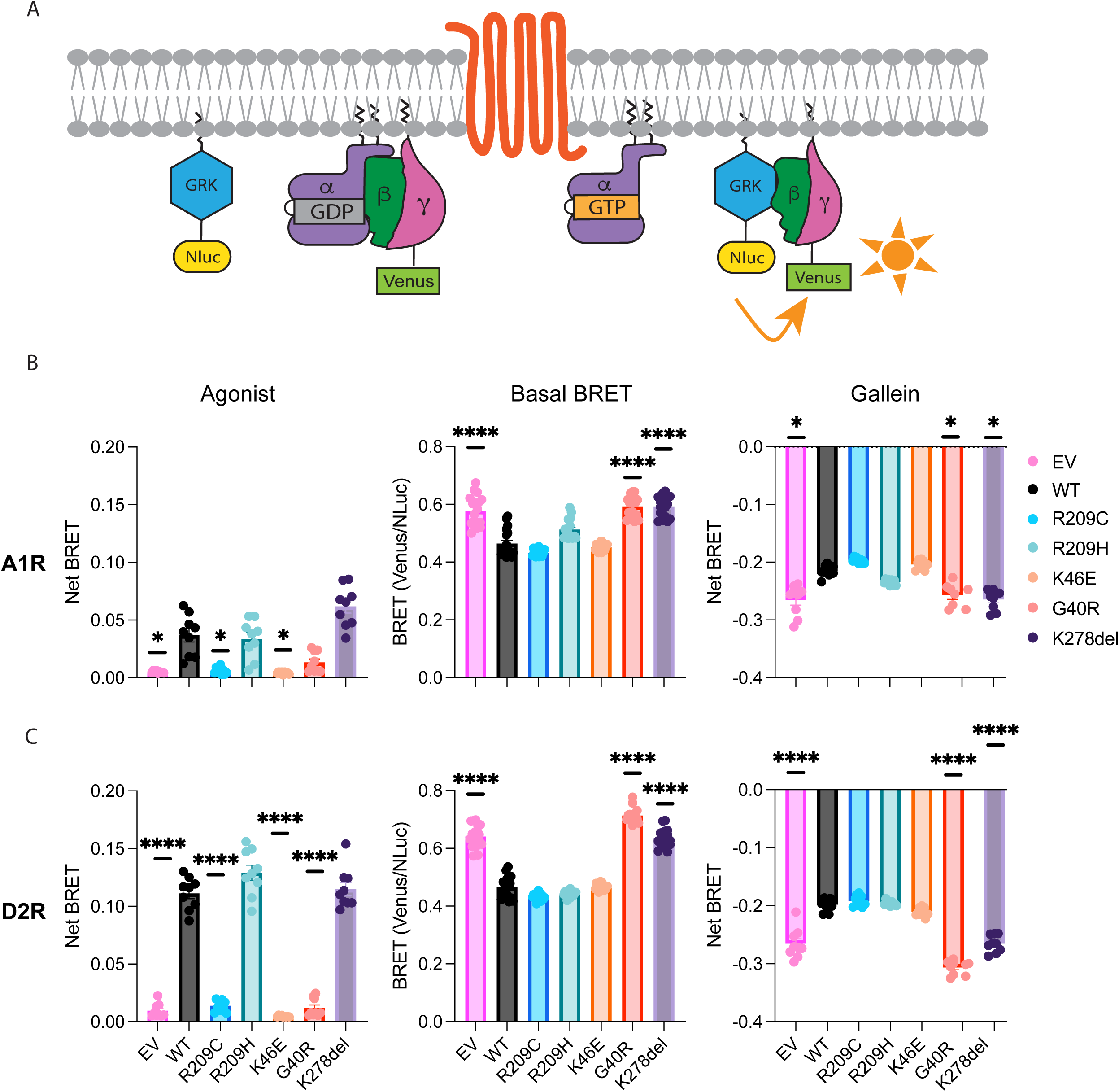
*GNAO1* R209 variants impair Gβγ-effector coupling. (A) Schematic of the Gβγ-GRK BRET assay. In the inactive heterotrimer state, Gα_o_ and Gβ_1_/Gγ_2_-Venus are in close proximity, producing a low Venus/Nluc BRET signal. Agonist-induced activation triggers heterotrimer dissociation, causing Gβ_1_/Gγ_2_-Venus to associate with masGRK3ct-Nluc-HA, increasing the BRET signal. (B–C) Net BRET in response to agonist (left), basal BRET in the absence of agonist (middle), and net BRET in response to the Gβγ inhibitor gallein (right) for the indicated Gα_o_ variants paired with the adenosine A1 receptor (A1R; B) or dopamine D2 receptor (D2R; C). K46E and G40R are non-functional benchmark controls; K278del, a full-codon deletion, is a positive control for poor Gα_o_ expression (Fig. S1). Bars represent mean ± SD with individual replicates shown. All experiments have at least 3 biological replicates with 3 technical replicates. Statistical analyses were performed on the mean of the technical replicates from each biological replicate. Statistical comparisons are to WT using one-way ANOVA with Dunnett’s post hoc test, *p < 0.05, ***p < 0.001, ****p < 0.0001.

While the agonist data identify a coupling defect, they do not reveal whether the R209 mutations alter steady state heterotrimer assembly or constitutive Gβγ release, or whether any such effect depends on the receptor. Basal BRET in this assay, measured without agonist, reflects how much Gβγ is bound to GRK at rest and indicates heterotrimer expression: when Gα_o_ is abundant, most Gβγ is sequestered in the heterotrimer state and basal BRET is low, whereas without Gα_o_ (EV), Gβγ is free to bind GRK and basal BRET is higher. We measured basal BRET for each variant at A1R and D2R, pooling replicates from the agonist and gallein conditions recorded before drug addition. We expected poorly expressed variants (EV, G40R, K278del) to show high basal BRET, while well-expressed variants should resemble WT. As expected, WT showed lower basal BRET than EV, G40R, and K278del at both receptors (Fig. 3B and C, middle panel), consistent with the poor expression of these variants (Fig. S1). All other variants: R209C, R209H, and K46E were no different than WT at the A1 and D2 receptors (Fig. 3B and C, middle panel and Fig. S3A and B). These data raise the question of whether a receptor-independent intervention can override poor receptor coupling and reduced Gα_o_ expression to restore signaling, as gallein did in the TRUPATH experiments.

As shown in Fig. 2, gallein rescued signaling for both normally and poorly expressed Gα_o_ variants. We next applied gallein to cells expressing each variant at A1R and D2R and measured Gβγ– GRK association. EV, WT, and the poorly expressed variants served as controls. Gallein produced a sharp decrease in net BRET, consistent with its blocking of the Gβγ–GRK interaction. The largest decreases occurred for EV, G40R, and K278del at both receptors (Fig. 3B and C, rightmost panel). At A1R and D2R, R209C, R209H, and K46E all showed dissociation comparable to WT (Fig. 3B and C, rightmost panel and Fig. S3C and D). Overall, this activity pattern tracks heterotrimer expression. The variants that are poorly expressed or absent (EV, G40R, K278del) exhibited the highest basal BRET (Fig. 3B and C, middle panel), leaving a substantial Gβγ–GRK complex for gallein to disrupt and resulting in the largest net change. Together, these results show that gallein acts on Gβγ across two orthogonal assays and suggest it does so by promoting heterotrimer dissociation independently of receptor activation.

### *GNAO1*^R209H/+^ mice show altered intrinsic membrane properties in striatal SPNs

Gα_o_ is highly expressed in the striatum (https://proteinatlas.org), a basal ganglia structure critical for regulating motor behavior, the *GNAO1*^R209H/+^ mice display hyperkinesis, and direct pathway D1 SPNs are often considered to promote movement.^10,23,33,35,36^ This led us to investigate the effects of R209H mutation on the intrinsic membrane properties of striatal SPN subtypes. To address this, we performed whole-cell current-clamp recordings from SPNs in the dorsolateral striatum of WT and *GNAO1*^R209H/+^ littermates of both sexes. D1 SPNs were identified by expression of a *Drd1*-tdTomato reporter, and non-fluorescent neurons were classified as putative D2 SPNs.

Within D1 SPNs, resting membrane potential (RMP) showed no significant change between genotypes in either sex (Tables S4 and S5). Only D1 SPNs from male *GNAO1*^R209H/+^ mice exhibited significantly larger hyperpolarizing voltage deflections and input resistance than their WT littermates but no significant difference in membrane resistance (Fig. 4A). In contrast, D1 SPNs from female *GNAO1*^R209H/+^ mice were indistinguishable from WT across all measured parameters, including RMP, voltage deflection, input resistance, and membrane resistance (Fig. 4B). These findings provide a potential electrophysiological explanation for the hyperlocomotion observed in male but not in female *GNAO1*^R209H/+^ mice and raise the question of whether the locomotor phenotype in females arises instead from intrinsic dysfunction within D2 SPNs.

**Figure 4.**
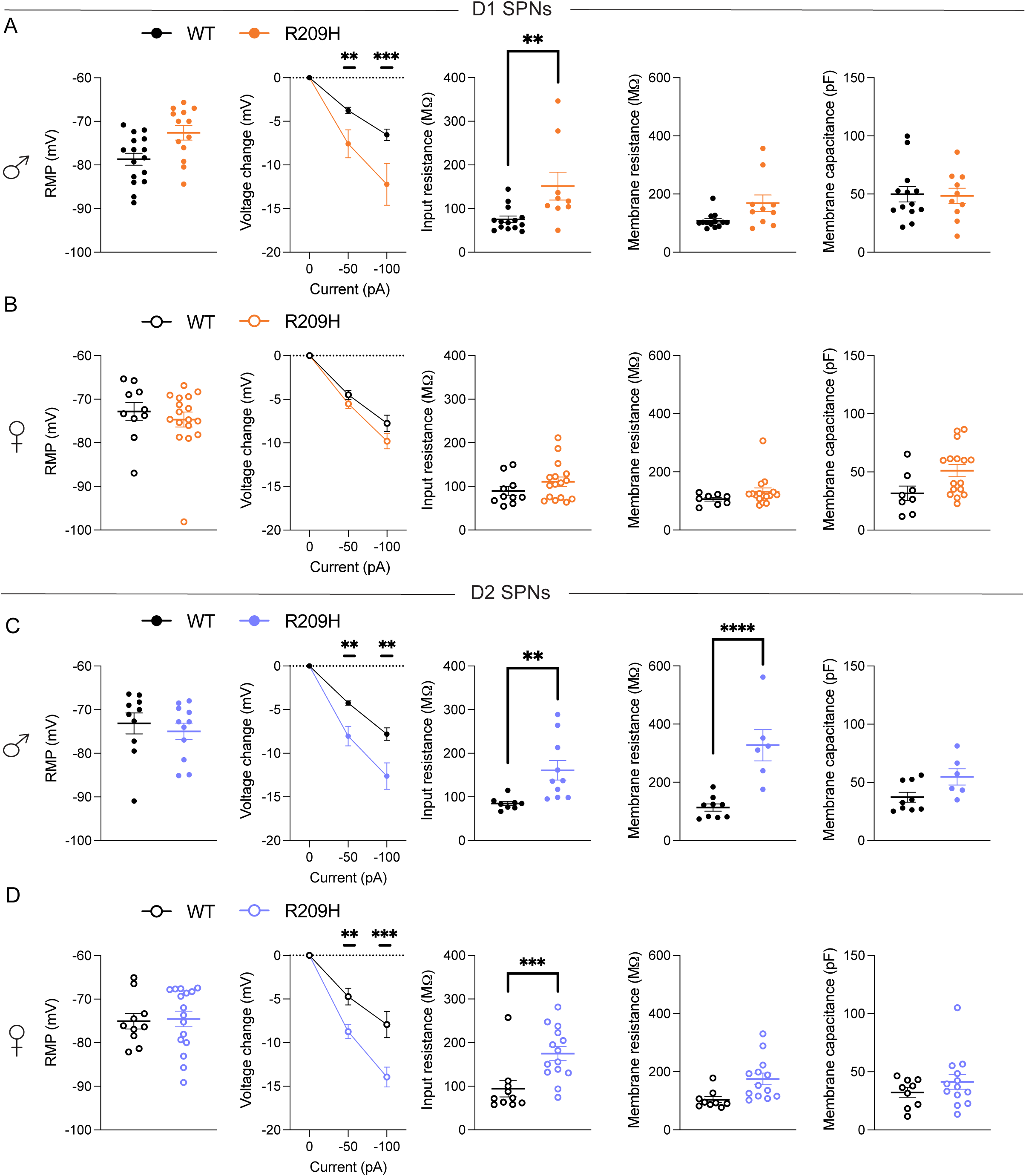
The Gα_o_ R209H mutation alters intrinsic membrane properties of striatal SPNs in a cell-type- and sex-specific manner. (A–B) Intrinsic membrane properties of D1 SPNs recorded in the dorsolateral striatum of WT male (A, black closed circles, n = 16 cells, N = 7), *GNAO1*^R209H/+^ male (A, orange closed circles, n = 13 cells, N = 9), WT female (B, black open circles, n = 10 cells, N = 6) and *GNAO1*^R209H/+^ female (B, orange open circles, n = 17 cells, N = 6) mice. From left to right: resting membrane potential (RMP), voltage change in response to hyperpolarizing current injection (−100 pA, −50 pA), input resistance, membrane resistance, and membrane capacitance. (C–D) Intrinsic membrane properties of putative D2 SPNs from WT male (C, black closed circles, n = 10 cells, N = 6), *GNAO1*^R209H/+^ male (C, blue closed circles, n = 11 cells, N = 8), WT female (D, black open circles, n = 10 cells, N = 5) and *GNAO1*^R209H/+^ female (D, blue open circles, n = 16 cells, N = 9) mice. Voltage change and input resistance panels reflect a subset of the above: male D1 WT 14/5, R209H 9/8; female D1 WT 10/6, R209H 16/6; male D2 WT 8/5, R209H 10/7; female D2 WT 10/5, R209H 14/9 (cells/mice). Membrane resistance and capacitance panels: male D1 WT 13/6, R209H 10/7; female D1 WT 8/5, R209H 16/6; male D2 WT 9/5, R209H 6/6; female D2 WT 9/4, R209H 13/8. Data points represent individual cells; bars represent mean ± SEM. Statistics: linear mixed-effects models with mouse identity as a random intercept to account for multiple cells recorded from the same animal (genotype × sex × cell-type fixed effects, × current step for voltage change; input resistance, membrane resistance, and membrane capacitance modeled on a log scale); genotype contrasts Holm-corrected within each sex × cell-type group. Complete statistics for the omnibus term and every comparison are in Tables S4 and S5, respectively. **p < 0.01, ***p < 0.001, ****p < 0.0001.

We next examined whether the R209H mutation also altered D2 SPNs’ membrane properties. In both male and female *GNAO1*^R209H/+^ mice, D2 SPNs showed no significant changes in RMP, yet significantly greater hyperpolarizing voltage deflections and increased input resistance compared to WT (Fig. 4C and D). Membrane resistance was elevated in the D2 SPNs of male *GNAO1*^R209H/+^ mice but not in females (Fig. 4C and D). Neither membrane capacitance nor access resistance differed by genotype, sex, or cell type (Fig. 4A-D, rightmost panel), arguing against differences in cell size or recording quality as confounds. The elevated input and membrane resistances reflect reduced functional ion channel conductance and indicate decreased synaptic input is required to produce a voltage response, consistent with larger voltage deflections. However, the present experiments do not identify the affected conductances and cannot rule out contributions from altered neuronal morphology.

### The R209H mutation increases cellular excitability in both direct and indirect pathway SPNs in a sex-specific manner

Since pharmacologic normalization of D1-SPN activity can alleviate hyperlocomotion,^10,37^ we theorized that increased excitability of D1 SPNs may underlie the hyperlocomotor phenotype. We tested the excitability of D1 SPNs by performing whole-cell current-clamp recordings across a range of current-injection steps (Fig. 5A). WT littermates acted as controls, and rheobase was used as a readout of the excitability threshold. Male *GNAO1*^R209H/+^ mice showed a significant increase in D1 SPN action potentials elicited and a lower rheobase relative to WT controls (Fig. 5B), whereas female *GNAO1*^R209H/+^ D1 SPNs did not significantly differ from WT in either measure (Fig. 5C). The elevated input resistance observed in male D1 SPNs (Fig. 4A) provide a mechanistic basis for this increased excitability: a higher input resistance amplifies the voltage response per unit of injected current, reducing the current required to drive spiking, and increasing gain across the frequency-current relationship. There was a significant genotype × current-step interaction and a significant genotype × cell-type × current-step interaction, indicating that the effect of genotype on this I-V relationship differs between D1 and D2 SPNs, consistent with previous work demonstrating that D2 SPNs are more excitable than D1 SPNs.^38^

**Figure 5.**
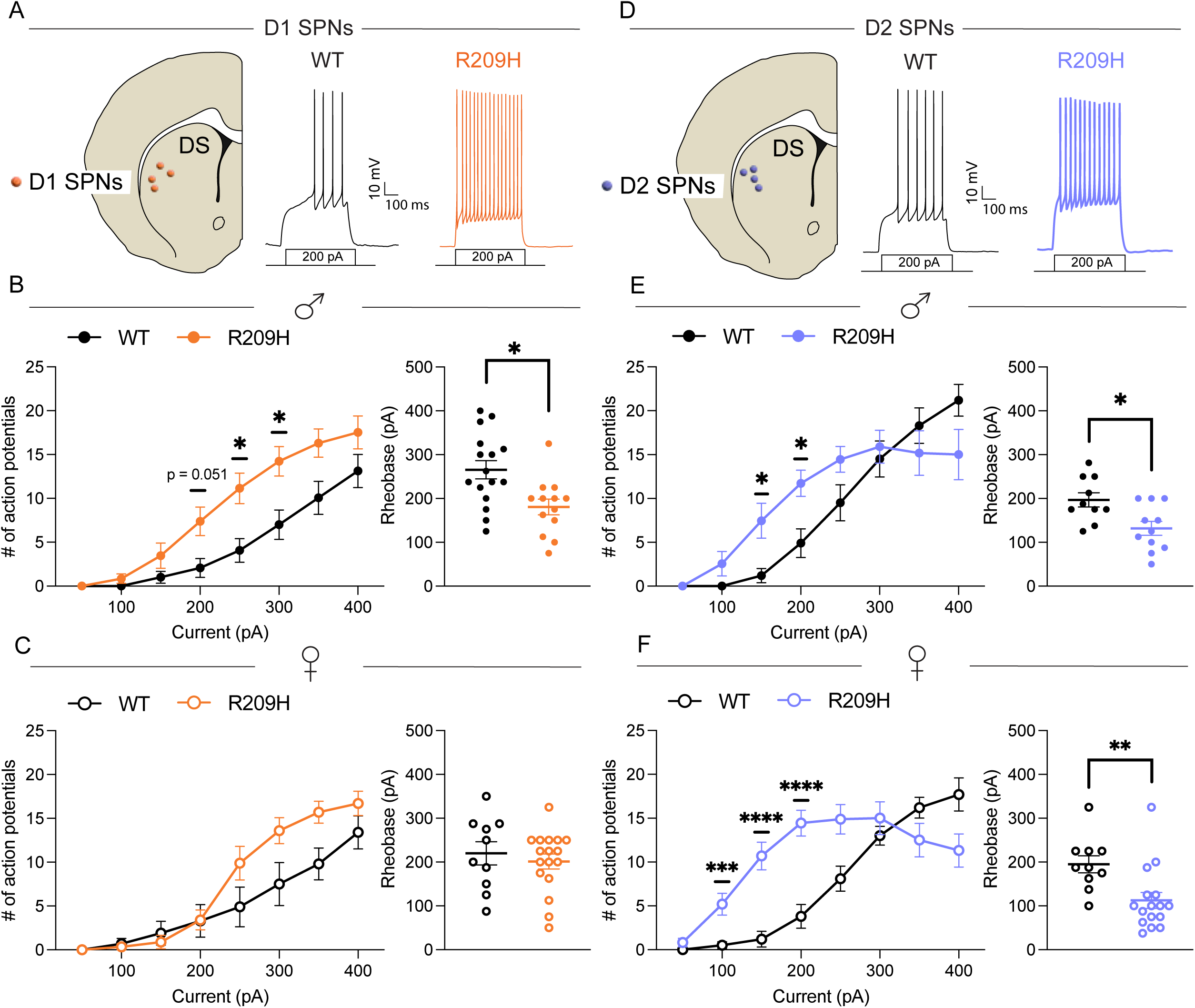
The Gα_o_ R209H mutation produces sex- and cell-type-specific alterations in striatal SPN excitability. (A and D) Schematic diagrams of the mouse dorsal striatum indicating the recorded cell populations: D1 SPNs (A) and D2 SPNs (D). Representative whole-cell current-clamp traces are shown for WT (black) and *GNAO1*^R209H/+^ (orange and blue) neurons in response to a 200 pA current injection step (scale bars: 10 mV, 100 ms). (B) Input-output curves (left) and rheobase (right) for D1 SPNs recorded from male WT littermate controls (black, n = 16 cells, N = 7 mice) and male *GNAO1*^R209H/+^ (orange, n = 13 cells, N = 9) mice. (C) Input-output curves and rheobase for D1 SPNs from female WT (open black circles, n = 10 cells, N = 6) and female *GNAO1*^R209H/+^ (open orange circles, n = 17 cells, N = 6) mice. (E) Input-output curves and rheobase for D2 SPNs from male WT (black, n = 10 cells, N = 6) and male *GNAO1*^R209H/+^ (blue, n = 11 cells, N = 8) mice. (F) Input-output curves and rheobase for D2 SPNs from female WT (open black circles, n = 10 cells, N = 5) and female *GNAO1*^R209H/+^ (open blue circles, n = 16 cells, N = 9) mice. Data are presented as mean ± SEM. Statistics: input-output curves used a nested repeated-measures model (current step within cell within mouse; mouse random intercept; genotype × sex × cell-type × current-step fixed effects); rheobase used a log-scale linear mixed-effects model (mouse random intercept). Genotype contrasts were Holm-corrected within each sex × cell-type curve (8 current steps) or stratum (rheobase). Complete statistics for the omnibus term and every comparison are in Tables S6 and S7, respectively. *p < 0.05, **p < 0.01, ***p < 0.001, ****p < 0.0001.

Classically, indirect pathway D2 SPNs are thought to counterbalance direct pathway activity by inhibiting thalamic activity, thereby suppressing unwanted movement.^11–13,39^ Thus, the hyperlocomotion in females could result from hypoexcitability of D2 SPNs. Male *GNAO1*^R209H/+^ D2 SPNs showed lowered rheobase and hyperexcitability at intermediate current injections (150–200 pA) but decreased firing at higher current steps (350–400 pA). This biphasic response could be interpreted as depolarization block due to sodium channel inactivation (Fig. 5D, E).^40^ Female *GNAO1*^R209H/+^ D2 SPNs displayed a qualitatively similar profile, with lowered rheobase, hyperexcitability at lower current steps (100–200 pA), and suppressed firing at higher steps (400 pA, Fig. 5F). This high-current downturn in D2 SPNs of both sexes is directionally consistent with a depolarization-block-like process but did not reach statistical significance. These patterns are consistent with the elevated input resistance shared by male and female D2 SPNs (Fig. 4C, D). This combination likely renders these cells more prone to depolarization but less capable of sustained activity at high-current steps. Together, these results indicate that R209H increases SPN excitability through elevated input resistance and reduced rheobase, with a firing phenotype that differs significantly in its current-dependent shape between D1 and D2 SPNs. A sex-dependent component is most clearly established in D1 SPNs, where the excitability phenotype is present in males and absent in females, whereas D2 SPNs show hyperexcitability in both sexes. This dissociation suggests divergent contributions of direct and indirect pathway activity to the motor phenotype across sexes.

### Intra-DLS infusion of gallein and selective adenosine receptor antagonists reduce hyperlocomotion in *GNAO1*^R209H/+^ mice

The dorsolateral striatum (DLS) is a central hub for motor control, and restoration of WT *GNAO1* in the striatum is sufficient to reduce hyperlocomotion.^41^ Further, we find that SPNs in this region are hyperexcitable in *GNAO1*^R209H/+^ mice (Fig. 5). Therefore, we next investigated whether local pharmacological modulation of the DLS is sufficient to reduce the hyperlocomotor phenotype. To test this, we bilaterally cannulated mice targeting the DLS and infused gallein or vehicle using a within-subject design (Fig. S4A, B). Only female *GNAO1*^R209H/+^ mice showed characteristic hyperactivity within the vehicle condition, which was significantly reduced to WT levels following intra-DLS gallein (Fig. S4D). Male *GNAO1*^R209H/+^ mice hyperactivity within the vehicle condition just missed significance with a p - value of 0.053 (Fig. S4C). These findings support a contribution of DLS Gβγ signaling to GNAO1-related hyperlocomotion. However, because gallein acts broadly across all cell types in the region, we sought to identify a specific receptor locus that may underlie this effect.

Prior work has shown that A1 and A2A receptors differentially regulate locomotion and locomotion-associated PKA signaling in direct- and indirect-pathway SPNs.^23^ We therefore compared local DLS infusion of the A1R antagonist DPCPX and the A2AR antagonist KW-6002 to determine whether either receptor contributed to the altered locomotor response in *GNAO1*^R209H/+^ mice. We used the same cannulated cohort and sequentially infused KW-6002 and DPCPX into the DLS, with open field testing following each infusion (Fig. S4A, B). In male *GNAO1*^R209H/+^ mice, DPCPX appears to reduce hyperlocomotion to wild-type levels, whereas KW-6002 had no effect in either sex (Fig. S4E, F). DPCPX had no effect on female *GNAO1*^R209H/+^ locomotion (Fig. S4F). Taken together, these data suggest Gβγ and A1R signaling in the DLS as potential therapeutic targets for GNAO1-related hyperkinesis.

### Systemic administration of two small molecules reduces hyperlocomotion in *GNAO1*^R209H/+^ mice

Previous work demonstrated that *GNAO1*^R209H/+^ mice exhibit robust hyperlocomotion (open-field locomotor activity; Fig. 6A), providing a translatable readout for assessing the efficacy of small-molecule therapeutics in vivo.^10^ We next sought to assess the therapeutic potential of systemic modulation of Gβγ and adenosine signaling. First, we systemically administered gallein to male and female *GNAO1*^R209H/+^ and WT littermates in a crossover intraperitoneal (i.p.) dosing regimen, in which each animal received vehicle and gallein across separate testing days, followed by open-field assessment of locomotor activity (Fig. 6B). Gallein significantly reduced hyperlocomotion in both sexes in *GNAO1*^R209H/+^ mice. While male *GNAO1*^R209H/+^ mice were still more active than WT (Fig. 6C), female mice exhibited a pronounced reduction in activity to WT levels (Fig. 6D). These findings demonstrate that pharmacologic modulation of Gβγ signaling is sufficient to rescue the motor phenotype in *GNAO1*^R209H/+^ mice.

**Figure 6.**
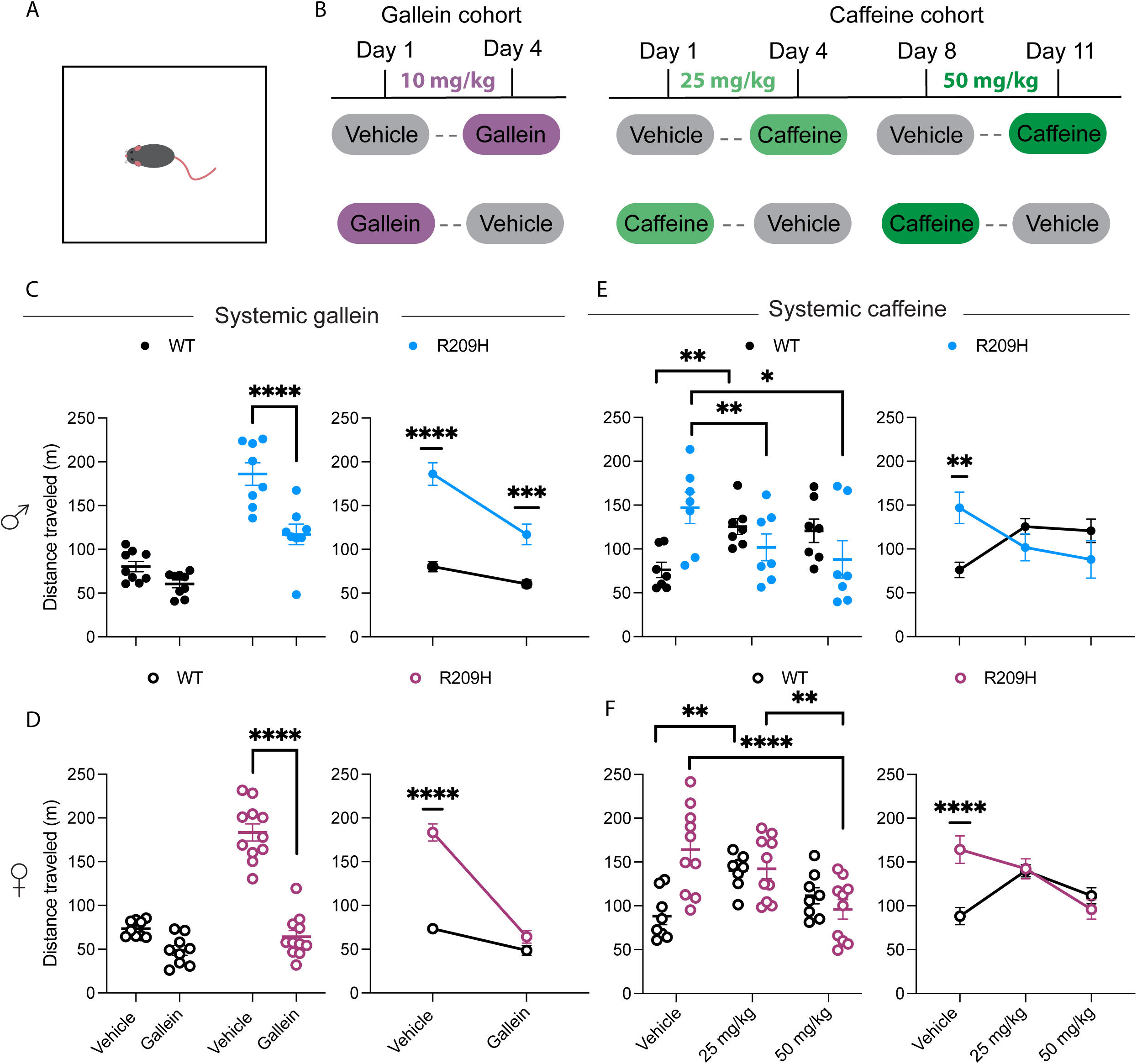
Systemic administration of two small molecules reduces hyperlocomotion in *GNAO1*^R209H/+^ mice in the open field test. (A) Schematic of the open field arena. (B) Experimental timeline for the crossover dosing regimen. Each animal received both vehicle and drug treatments on separate testing days, with an open-field assessment following each injection. Effect of systemic gallein on locomotor activity in (C) male WT and *GNAO1*^R209H/+^ (WT, n = 9; R209H, n = 8) and (D) female WT and *GNAO1*^R209H/+^ (WT, n = 9; R209H, n = 11) mice. Left panels show individual data points and show within-genotype comparisons; right panels show paired means ± SEM with between-genotype comparisons. Effect of systemic caffeine (25 or 50 mg/kg) on locomotor activity in (E) male WT and *GNAO1*^R209H/+^ (WT, n = 7; R209H, n = 7) and (F) female WT and *GNAO1*^R209H/+^ (WT, n = 8; R209H, n = 10) mice. Gallein data were analyzed using a two-way repeated-measures ANOVA with Fisher’s LSD post hoc test. Caffeine data were analyzed with a two-way repeated-measures ANOVA with Tukey’s post hoc test. F statistics and post hoc test p-values can be found in Tables S8 and S9, respectively. *p < 0.05, **p < 0.01, ***p < 0.001, ****p < 0.0001.

Caffeine has been proposed as a therapeutic for GNAO1-related disorders, although its efficacy has only been demonstrated in non-mammalian systems.^24^ To determine whether caffeine reduces hyperlocomotion in an mammalian model system, we used the same crossover i.p. dosing regimen; each animal received vehicle, 25 mg/kg caffeine, or 50 mg/kg caffeine across separate testing days (Fig. 6B). In *GNAO1*^R209H/+^ males, both the 25 and 50 mg/kg doses significantly reduced hyperlocomotion relative to vehicle (Fig. 6E). In females, only the 50 mg/kg dose significantly reduced locomotion toward WT levels (Fig. 6F). 25 mg/kg caffeine increased locomotion in WT mice of both sexes consistent with prior reports.^42^ Together, these results demonstrate that caffeine normalizes locomotion in *GNAO1*^R209H/+^ mice, and does so in a dose- and sex-dependent manner. These results support the therapeutic potential of caffeine, while also highlighting dose and sex as critical variables, and may help to explain the differential effects of DLS infusion of DPCPX.

## Discussion

Here, we integrated human genetic analysis with cellular assays to define mechanistic classes of *GNAO1* mutations and *in vivo* approaches to evaluate therapeutic strategies. Our analysis of patient data revealed that mutations that impair Gβγ release are significantly associated with NEDIM. Electrophysiological studies revealed sex- and cell-type–specific alterations in spiny projection neurons. Using a mouse model of NEDIM, we showed that gallein and caffeine significantly reduce hyperlocomotion in the open field test. Together, these findings implicate dysfunction of the dorsolateral striatum in GNAO1-related disorders and highlight the potential of two pharmacological agents to restore proper movement.

Our data support a model in which Gα_o_ acts as a sink for Gβγ, capable of rapidly switching signaling on and off. ^26^ Gβγ regulates adenylyl cyclase as well as ion channels, including G protein-coupled inwardly rectifying potassium channels, and components of the SNARE complex.^43–46^ Gα_o_ itself shapes cAMP production; however, unlike other Gα_i/o_ family members, it does not directly inhibit adenylyl cyclase.^47^ This is instead attributed to Gα_o_ control of free Gβγ, which in turn binds and modulates adenylyl cyclase activity.^43^ Recently, Rap1GAP1a was identified as a candidate Gα_o_ effector,^48^ and that same analysis showed that the R209H mutation impairs Rap1GAP1a–Gα_o_ interaction. Even so, the biophysical properties of Gα_o_ are consistent with a primary role as a regulator of Gβγ availability: mutations at R209 that constitutively sequester Gβγ are sufficient to drive hyperkinesis possibly by altering neuronal excitability, although alterations in neurotransmitter release may also contribute to this phenotype. Further, pharmacological modulation of Gβγ signaling with gallein restores normal motor behavior. These data place Gβγ dysregulation at the mechanistic core of GNAO1-related NEDIM.

If Gα_o_ functions primarily as a sink for Gβγ, then liberating Gβγ pharmacologically should be a therapeutically viable option. Gallein is a small molecule identified in a screen for compounds that directly bind Gβγ.^25^ It associates and dissociates slowly and disrupts Gβγ interactions with downstream effectors including PLCβ2, PLCβ3, PI3K, and GRK.^25,49^ Although gallein has been shown to disrupt Gβγ binding to these effectors, its effect on Gβγ–Gα_o_ interaction has not previously been tested.^25^ This is an important gap, given that increased subunit-subunit interaction likely accounts for the NEDIM phenotype. Here we show that gallein alters the Gβγ– Gα_o_ interaction. We show further that gallein reduces hyperlocomotion, following both systemic administration and direct infusion into the dorsolateral striatum, selectively in *GNAO1*^R209H/+^ but not in wildtype mice. These findings are consistent with prior work showing that gallein is well tolerated systemically and, following bilateral infusion into the nucleus accumbens, attenuates amphetamine-induced but not basal locomotor activity.^50^ Gallein has also demonstrated efficacy in mouse models of heart failure, pain, and inflammation,^51–54^ underscoring its therapeutic breadth. Together, these data position gallein as a promising therapeutic candidate for GNAO1-related disorders and one that is mechanistically distinct from, and potentially complementary to, therapies like caffeine that target receptors.

Our caffeine findings closely parallel those reported in a conditional *GNAO1*^G203R/+^ model.^55^ Brunori et al. found that 20 mg/kg caffeine increased locomotion in control mice but reduced hyperactivity and improved rotarod performance when G203R expression was restricted to striatal neurons. Together with our findings in *GNAO*1^R209H/+^ mice, these results provide convergent evidence across two pathogenic variants that caffeine can improve GNAO1-related motor dysfunction and further implicate the striatum as an important site of action. The receptor-level mechanism, however, remains unclear. Brunori et al. interpreted their findings in the context of altered A2AR signaling in indirect-pathway SPNs, whereas our intra-DLS experiments point to the involvement of A1R antagonism. This difference may reflect variant-specific signaling, restriction of G203R expression to striatal neurons, or differences between systemic caffeine and local receptor-selective antagonism. Thus, the behavioral efficacy of caffeine is supported across models, but the receptor subtype and circuit mechanisms remain unresolved.

An unusual feature of GNAO1-related disorder is its female predominance (Fig. 1D), in contrast to other neurodevelopmental disorders, including autism spectrum disorder and attention-deficit/hyperactivity disorder.^27–29^ This pattern is incongruent with the Female Protective Effect, a liability threshold model, that posits that biological females require a greater burden of genetic and environmental risk factors to produce neurodevelopmental phenotypes.^56^ Unlike Rett syndrome, where female predominance is explained by male lethality stemming from the X-linked location of *MECP2*, *GNAO1* is an autosomal gene, ruling out X-chromosome inactivation as a contributing mechanism.^31,57,58^ Our electrophysiological findings offer one explanation for the observed sex differences: the R209H mutation produces distinct alterations in SPN excitability in males and females, with D1 SPNs hyperexcitable in males and D2 SPNs exhibiting a biphasic change in excitability in both sexes. Interestingly, selective deletion of *GNAO1* in D1-MSNs causes a similar change in their excitability, suggesting that any impairment in Gα_o_ can induce dysregulation of their activity.^34^ The rescue achieved by gallein likely reflects its ability to act intracellularly at the level of Gβγ, rather than at the receptor level as seen with A1R antagonism. Whether the sex-specific divergence in SPN vulnerability reflects a mechanism that renders females more susceptible to GNAO1-related disorder, or instead underlies differences in how the disorder manifests between the sexes, remains an important question for future study.

A sex-specific SPN divergence has been reported previously in another model of neurodevelopmental disorder. In mice carrying the 16p11.2 hemi-deletion, a copy-number variant associated with autism and intellectual disability, D2 SPN hemi-deletion produces male-specific hyperactivity, whereas cortical hemi-deletion increases hyperactivity in both sexes.^59^ Our data extend this principle and offer a mechanistic resolution via equifinality: both male and female *GNAO1*^R209H/+^ mice exhibit hyperlocomotion, but males show disruptions in both direct and indirect pathways, whereas females only have altered indirect-pathway function. This distinction may have direct therapeutic implications. The globus pallidus internal is an effective target for deep brain stimulation (DBS) in GNAO1-related disorder,^15–17,60,61^ and an ongoing clinical trial (NCT06912841) is examining the effectiveness of DBS in childhood-onset hyperkinetic disorders, including GNAO1-related disorder. The globus pallidus internal is where direct and indirect pathways converge, likely accounting for its importance as a DBS target. However, no published data stratify DBS outcomes by sex, leaving open the question of whether sex-specific circuit differences translate into sex-specific treatment responses in human patients.

Together, our findings provide a mechanistic and circuit-level framework for understanding and treating GNAO1-related hyperkinetic movement disorder. We demonstrated that Gβγ sequestering mutations are strongly and consistently associated with NEDIM across a large patient cohort. These mechanistic classifications carry direct therapeutic implications: patients with haploinsufficient mutations, in whom Gα_o_ protein is absent or mislocalized, may not benefit from approaches that increase Gβγ interactions with effectors. In contrast, those with Gβγ-sequestering variants such as R209H represent a mechanistically defined population for whom increasing free Gβγ is likely to be beneficial. This work illustrates how integrating cellular assays, patient cohort data, and in vivo circuit analysis can generate a coherent, clinically actionable framework for a rare neurodevelopmental disorder.

## Materials and Methods

### Human data

All patient data were collected from previously published reports except for patients from the United States *GNAO1*-related Disorder Natural History Study, as denoted by the United States GNAO1-related Disorder Natural History Study (trials.gov identifier NCT04950946). These data were obtained with permission from clinicians and patients and de-identified. Patients lacking sex annotation were excluded from analyses. A full list of patients with corresponding citations is provided in Table S1. Variants were classified as haploinsufficient or as sequestering Gβγ based on published BRET and immunoblotting studies, with citations for each mutation provided in Table S2. Analysis code is available in a public repository (see Data availability).

### Cell lines

All TRUPATH cellular assays were performed using HEK293T/17 (ATCC; CRL-11268) cells maintained at 37 °C in Dulbecco’s Modified Eagle Medium (DMEM) (Corning; 10-017-CV) supplemented with 10% FBS (GIBCO; A3382001), 100 I.U./mL penicillin, and 100 mg/mL streptomycin. All other cellular assays were performed using HEK293T/17 cells maintained at 37 °C in DMEM supplemented with 10% FBS and 1% MEM Non-Essential Amino Acids Solution (Thermo Fisher; 11140050).

### Mutagenesis and plasmid construction.^62^

The only newly generated constructs were K278del in the pcDNA5 with Rluc8 and pcDNA3.1 backbones. Primestar Max (Takara Bio; R045A) mutagenesis was performed using primer pairs, each containing 5 bp 5’ and 20-30 bp 3’ to the relevant codon. The oligo sequences (5’ - 3’) are as follows. Gα_o_ K278del fwd: TTGGCGAGATCAAGAAGTCACCTTTGACCATCTGCTTTC. Gα_o_ K278del rev: TTCTTGATCTCGCCAAAGAGATCTTTCTTGTTGAGGAAG. The full plasmid map for K278del in the pcDNA5 with Rluc8 is in File S1, and the plasmid map for K278del in the pcDNA3.1 backbone is in File S2. All other variant constructs can be found in ^9^.

### TRUPATH BRET

The following method was adapted from ^32^. Cells were co-transfected with a 1:1:1:1 ratio of plasmids containing the human dopamine 2 receptor (*DRD2*) or adenosine 1 receptor (*ADORA1*), Gα_o_ (wild-type and mutant constructs) fused to *Renilla* luciferase 8 (Rluc8), Gγ_8_ fused to GFP2, and Gβ_3_, carried out using TransIT-2020 (Mirus; MIR 5400) transfection reagent following manufacturer guidelines. The next day, cells were plated into poly-L-lysine-coated 96-well white clear-bottom plates (Corning; 3903) at a density of ∼60,000 cells/well in DMEM supplemented with 1% dialyzed FBS, 100 I.U./mL penicillin, and 100 mg/mL streptomycin. The following day, cells were washed with assay buffer (Hank’s Balanced Salt Solution containing 20 mM (4-(2-hydroxyethyl)−1-piperazineethanesulfonic acid) (HEPES) pH 7.4, 0.1% bovine serum albumin, and 0.01% ascorbic acid, then incubated with 60 μL assay buffer containing 7.5 μM coelentrazine 400a (Nanolight; 340). After 10 min in the dark, 30 μL of agonist (3X concentration) in drug buffer was added to each well. After another 10 min in the dark, the plates were serially read four times using a ClarioSTAR FSX plate reader with 395 nm (Rluc8-coelenterazine 400a) and 510 nm (GFP2) filters for 1 sec per well. Measurements from the final read were used for all analyses. The BRET ratio was calculated as the ratio between acceptor (GFP) and donor (Rluc8) emission values. Net BRET values were determined by subtracting the BRET ratio from cells treated with vehicle from the entire dataset. Net BRET data for each dose-response curve were fit using the (inhibitor) vs. response (three parameter) function in GraphPad Prism (GraphPad Software Inc., San Diego, CA). All experiments were run with at least three biological replicates, each with two technical replicates.

### Gβγ-GRK BRET

The following method was adapted from ^63^. Approximately 40,000 cells per well were distributed in 96-well flat-bottom white microplates (Corning; 3903) in the above media supplemented with 0.1% Matrigel. The cells were then transfected with plasmids encoding the human dopamine D2 receptor (*DRD2*) or adenosine A1 receptor (*ADORA1*), as well as masGRK3ct-Nluc (comprised of the C-terminal 141 amino acids of GRK3, an N-terminal peptide directing myristoylation, the HA epitope tag and Nano luciferase), human Gγ_2_ fused to residues 1-155 Venus (Venus 1-155-Gγ_2_), human Gβ_1_ fused to Venus 156-239 (Venus 156-239-Gβ_1_), and human Gα_o_ or mutant constructs described above in a 1:1:1:1:2 ratio, carried out using Lipofectamine LTX Reagent with PLUS (Thermo Fisher; 15338100). After 24 h, the cells were washed once with BRET buffer (Dulbecco’s Phosphate-Buffered Saline + 0.5 mM MgCl_2_ and 0.1% glucose) and resuspended in 50 μL BRET buffer. Fifty μL of the Nluc substrate furimazine (Promega; N1120) in BRET buffer was added per well, as directed by the manufacturer, followed by 50 μL (4X concentration) of 100 μM dopamine (Millipore Sigma; H8502) or 50 μM adenosine (Tocris Biosciences; 3624). The plates were read using a PHERAstar FSX microplate reader (BMG Labtech) equipped with two emission photomultiplier tubes, enabling simultaneous detection of two emissions at the highest possible resolution of 10 msec per data point. All measurements were performed at 37 °C. The BRET signal was determined by calculating the ratio of the light emitted by the Venus-Gβ_1_γ_2_ (535 nm ± 30 nm) over the light emitted by the masGRK3ct-Nluc-HA (475 nm ± 30 nm). The average baseline value (basal BRET ratio, with no Gα transfected) recorded prior to agonist stimulation was subtracted from the experimental BRET signal values. All experiments were run with at least three biological replicates, each with three technical replicates except for G40R, which had one technical replicate excluded due to an overgrown well.

### Western blots

HEK293T/17 cells were transfected with 200 ng pcDNA3.1 Gα_o_ wild-type and mutants and grown at 37 ^°^C in Dulbecco’s Modified Eagle Medium (DMEM, Corning; 10-013-CV) supplemented with 10% fetal bovine serum (Sigma-Aldrich; F2442). After 48 h, cells were lysed by sonication (Qsonica; Q500) at room temperature (5 sec on, 10 sec off, 30% amplitude, repeated 3 times) in 1X phosphate-buffered saline pH 7, 150 mM NaCl) and 1% Triton-X-100, with Pierce Protease Inhibitor Tablets (Thermo Fisher; A32965). Twenty mg of protein lysates were separated by 4-15% acrylamide gel electrophoresis and then subjected to immunoblotting. Nitrocellulose membranes were blocked with 0.1% fish gelatin in 1X Tris-buffered saline with Tween-20 (TBST) and blotted with 1:500 Gα_o_ antibody A2 (Santa Cruz Biotechnologies, sc-13532) and 1:5,000 mouse IgG Fc binding protein conjugated to Horseradish Peroxidase (m-IgG Fc BP HRP, Santa Cruz Biotechnologies, sc-525409). Blots were imaged using Clarity Western ECL Substrate (Bio-Rad, 1705061) on a ChemiDoc Touch Imaging System (Bio-Rad). Gα_o_ expression was quantified via fluorescence intensity using ImageJ.

### Animals

All transgenic mice were bred in-house on a C57Bl/6J background. *GNAO1*^+/R209H^ mice were a gift from Dr. Richard Neubig^10^ and were bred as heterozygous pairs to generate wild-type littermate controls. *GNAO1*^+/R209H^ were crossed to heterozygous “D1-tdTomato” (B6.Cg-Tg(Drd1a-tdTomato)6Calak/J, Jackson Laboratory) mice for electrophysiological experiments. The mice were maintained on a 12-h light/dark schedule in a vivarium at 22°C and 40% humidity, with food and water available ad libitum. No statistical methods were used to predetermine sample size. A limitation of the present study is that an a priori sample size calculation was not performed. The cohort size was determined by the number of available mice, which was limited by breeding constraints.

All procedures were approved by the University of North Carolina Institutional Animal Care and Use Committee and were conducted in accordance with the Guidelines for the Care and Use of Laboratory Animals.

### Blinding

All experiments were conducted in a blinded manner, with assays performed and analyzed without knowledge of the genotype, and animals were randomized by cage before behavioral experiments.

### Open field test

Mice were injected i.p. and, 20 min later, were placed in the center of a white plexiglass box measuring 40 cm x 40 cm for 30 min and conducted during the light cycle. An overhead camera recorded the mouse’s activity, which was analyzed using BIOBSERVE (GmbH). Separate cohorts were used to evaluate systemic gallein and caffeine. A within-subject crossover design was used within each drug cohort. For cannula infusion OFT experiments, mice were infused with the drug over 3 min. Infusers were kept in the cannula for an additional two min. Immediately afterward, mice were placed in the center of the open field box and were run as described above. Drugs were dissolved in DMSO and 0.9% NaCl at a final concentration of 3% DMSO.

### Drugs

Drugs injected i.p. were dissolved as follows. Gallein (Tocris Biosciences; 3090) was dissolved in 4% DMSO and administered to mice at 10 mg/kg. Vehicle contained 0.9% NaCl in 4% DMSO. Caffeine vehicle, 25 mg/kg, and 50 mg/kg caffeine (Tocris Biosciences; 2793) were dissolved in 0.9% NaCl. These drugs were administered in a counterbalanced fashion for OFT testing, with the 50 mg/kg caffeine condition following the 25 mg/kg condition in the same cohort.

Cannula drugs were all dissolved in 0.9% NaCl at a concentration of 3% DMSO. The following drugs were used: gallein (2 mM), KW-6002/Istradefylline (1 μM, Tocris Biosciences; 5147), and DPCPX (100 nM, Tocris Biosciences; 0439). Animals were infused with 500 nL of drug/hemisphere at the above dosages.

### Stereotaxic surgeries

Mice were anesthetized using ketamine/xylazine (100 and 20 mg/kg, respectively) placed in a stereotaxic frame, and the skull surface was exposed. Burr holes were drilled into the skull. Cannulas were implanted bilaterally into the dorsal lateral striatum (bregma coordinates, 0° angle, anteroposterior: +1.0, mediolateral: +/- 1.8, dorsoventral: -3.1) and were secured to the skull using miniature screws (thread size 00–90 × 1/16, Antrin Miniature Specialties) and light-cured dental adhesive cement (Geristore A&B paste, DenMat) was applied over the skull surface and base of cannulas.

### Statistics

Patient data odds ratios were analyzed using Fisher’s exact test. For the BRET experiments, statistical comparisons are to WT using one-way ANOVA with Dunnett’s post hoc test. Statistical analyses were performed on the mean of the technical replicates from each biological replicate. Open field test analysis was performed using a two-way ANOVA. Significance levels were set at p < 0.05. All analyses, except for the patient data analysis, were performed using GraphPad Prism 10.0 (GraphPad Software).

### Electrophysiology data analysis

Electrophysiological recordings comprised 103 cells from 35 mice (WT and *GNAO1*^R209H/+^ littermates of both sexes). Because cells recorded from the same animal are not statistically independent, all comparisons were conducted using linear mixed-effects models with mouse identity as a random intercept, following established recommendations for nested experimental designs.^64–66^ For outcomes measured once per cell, including resting membrane potential, rheobase, input resistance, membrane resistance, and membrane capacitance, models were fit with genotype, sex, and cell type and their interactions as fixed effects and mouse identity as a random intercept (lme4 and lmerTest packages in R). Input resistance, membrane resistance, membrane capacitance, and rheobase were log-transformed prior to analysis given their right-skewed, strictly positive distributions; raw-scale models are reported alongside as sensitivity analyses (Tables S4-S7). Access resistance was modeled identically as a quality-control check for differential recording quality across genotype, sex, and cell type.

For outcomes measured repeatedly within each cell across a series of current-injection steps, including evoked action-potential count and hyperpolarization-induced voltage deflection, current step was included as an additional repeated within-cell fixed factor, with cell nested within mouse as the random-effects structure, fit using the nlme package with a variance function (varIdent) allowing residual variance to differ across current steps.

Main effects and interactions were tested with Type III F-tests using sum-to-zero contrast coding, appropriate given the unbalanced group sizes in this dataset (Satterthwaite degrees of freedom for lme4 models). Planned pairwise genotype (R209H vs WT) comparisons were performed within each sex × cell-type stratum and, for the current-step outcomes, within each current level using estimated marginal means (emmeans package), with Holm correction applied across the four sex × cell-type strata (or across current levels within each stratum, for the current-step outcomes). All statistical tests were two-tailed, with significance defined as P < 0.05.

Model fit was evaluated by inspecting residuals and checking each mixed model for a singular fit (using lme4’s isSingular). Complete diagnostic results and the cross-validation analyses performed for the affected outcomes are in Supplementary Tables S4 and S5 (Fig. 4 outcomes: input resistance) and in Supplementary Tables S6 and S7 (Fig. 5 outcomes: rheobase).

All analyses were performed in R (version 4.5.2) using lme4 (version 1.1-38), lmerTest (version 3.2-1), nlme (version 3.1-168), and emmeans (version 2.0.4). Analysis code is available in a public repository (see Data availability).

### Slice electrophysiology

Mice were anesthetized with isofluorane and perfused immediately with ice-cold artificial cerebrospinal fluid (aCSF), which contains 128 mM NaCl, 3 mM KCl, 1.25 mM NaH_2_PO_4_ pH 7.4, 10 mM D-glucose, 25 mM NaHCO_3_, 2.5 mM CaCl_2_ and 2 mM MgCl_2_ (saturated with 95% O_2_ and 5% CO_2_, 295–305 mOsm). Acute coronal slices for dorsal lateral striatum patch clamp recordings were cut using a vibratome in cold sucrose (254 mM) aCSF saturated with 95% O_2_ and 5% CO_2_. Slices were maintained in holding chambers with aCSF for 1 h at 32°C. For current-clamp recordings, 3-6 MΩ electrodes were filled with potassium-based internal: 130 mM potassium methyl sulfate, 10 mM HEPES, 10 mM KCl, 2 mM MgCl2, 0.4 mM EGTA, 2.5 mM MgATP, and 0.3 mM NaGTP (pH 7.2-7.3, 283-287 mOsm). Recordings were performed using a MultiClamp 700B and a Digidata 1440, with a 2 kHz filter and 10 kHz digitization. Data acquisition and analysis were performed using the pClamp software Clampex 11.2. The Clampex Membrane Test was used to acquire membrane capacitance (Cm), membrane resistance (Rm), and access resistance. Cells with greater than 35 MΩ access resistance were discarded. Each cell-level value is the average of two technical sweeps from the same cell. Input resistance was calculated using a -50 pA hyperpolarizing step. Resting membrane potential was collected from the Clampex Membrane Test. To measure the intrinsic membrane properties, whole-cell recordings were carried out in current-clamp mode at the resting membrane potential and spikes were induced by incremental increases of current injection (each step increase was 50 pA; range -100–400 pA, 500 ms pulse). The number of action potentials was automatically counted using Clampfit Event Detection. Rheobase was calculated from the average first current step to induce an action potential(s).

## Data Availability

Analysis code for the electrophysiological and patient-cohort analyses is openly available at https://github.com/Gaby-soden/A-mechanistic-classification-of-GNAO1-mutations-links-G-sequestration-to-hyperkinetic-disease-and (archived at 10.5281/zenodo.22104353). The underlying electrophysiological and patient-cohort data are not included in the repository.

## Acknowledgments

We thank Dr. Richard Neubig for providing the *GNAO*1^R209H/+^ mice, Dr. Alan Smrcka for his comments on the manuscript and Dr. Thomas Kash and Dr. Michael Kasten for reviewing the electrophysiological findings and figures and providing feedback. We also acknowledge assistance from the NC Translational and Clinical Sciences (NC TraCS) Institute, which is supported by the National Center for Advancing Translational Sciences (NCATS), National Institutes of Health, through Grant Award Number UM1TR004406.

## Supplemental information

Document S1. Figures S1–S4 with corresponding figure legends and supplemental references.

Table S1. Patient mutations with phenotype and citations.

Table S2. Variant mechanism with corresponding citation.

Table S3. Logistic regression of NEDIM status on mechanism and sex (full model with interaction).

Table S4. Omnibus Type III tests — Figure 4 (membrane properties) outcomes.

Table S5. Pairwise genotype contrasts — Figure 4 (membrane properties) outcomes.

Table S6. Omnibus Type III tests — Figure 5 (excitability) outcomes.

Table S7. Pairwise genotype contrasts — Figure 5 (excitability) outcomes.

Table S8. F statistics for Figure 6.

Table S9. Post hoc test p-values for Figure 6.

Table S10. F statistics for Figure S4.

Table S11. Post hoc test p-values for Figure S4.

Data S1. Prism file corresponding to Figure 1.

Data S2. Prism file corresponding to Figure 2 and S2.

Data S3. Prism file corresponding to Figure 3 and S3.

Data S4. Prism file corresponding to Figure 4 and 5.

Data S5. Prism file corresponding to Figure 6-gallein data.

Data S6. Prism file corresponding to Figure 6-caffeine data.

Data S7. Prism file corresponding to Figure S4.

Data S8. Electrophysiology raw data in Excel.

File S1. Plasmid map of Gα_o_ K278del with Rluc8 in pcDNA5 backbone.

File S2. Plasmid map of Gα_o_ K278del in pcDNA3.1 backbone.

