## Supplementary files for "A mechanistic classification of *GNAO1* mutations links Gβγ sequestration to hyperkinetic disease and treatment response"

A

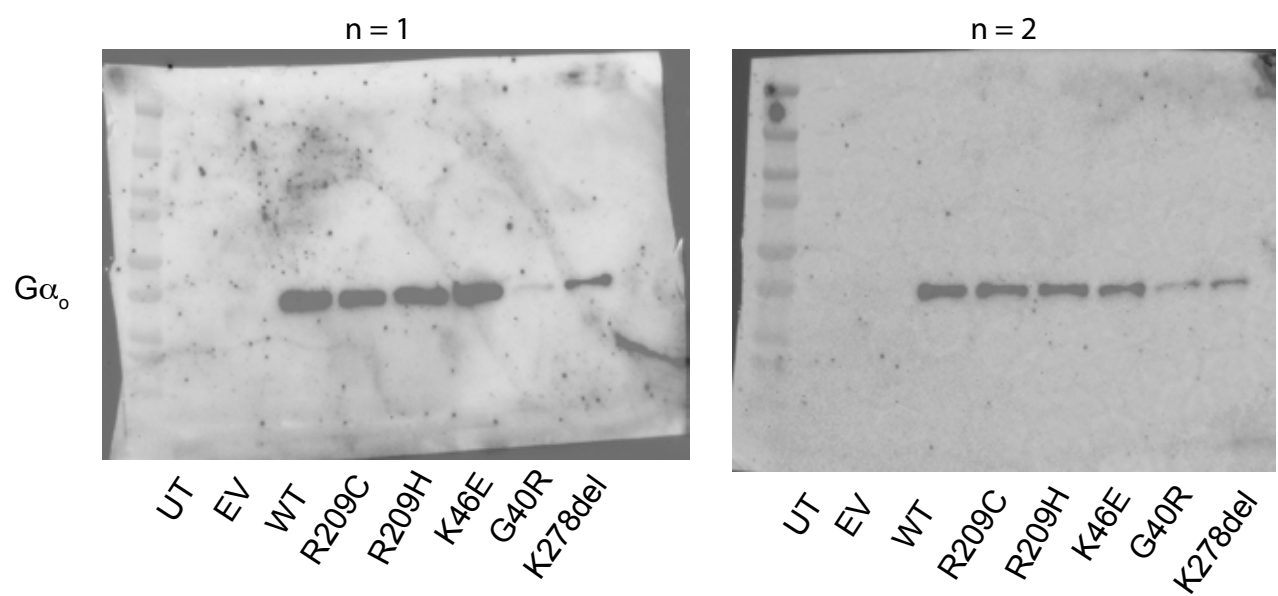

B

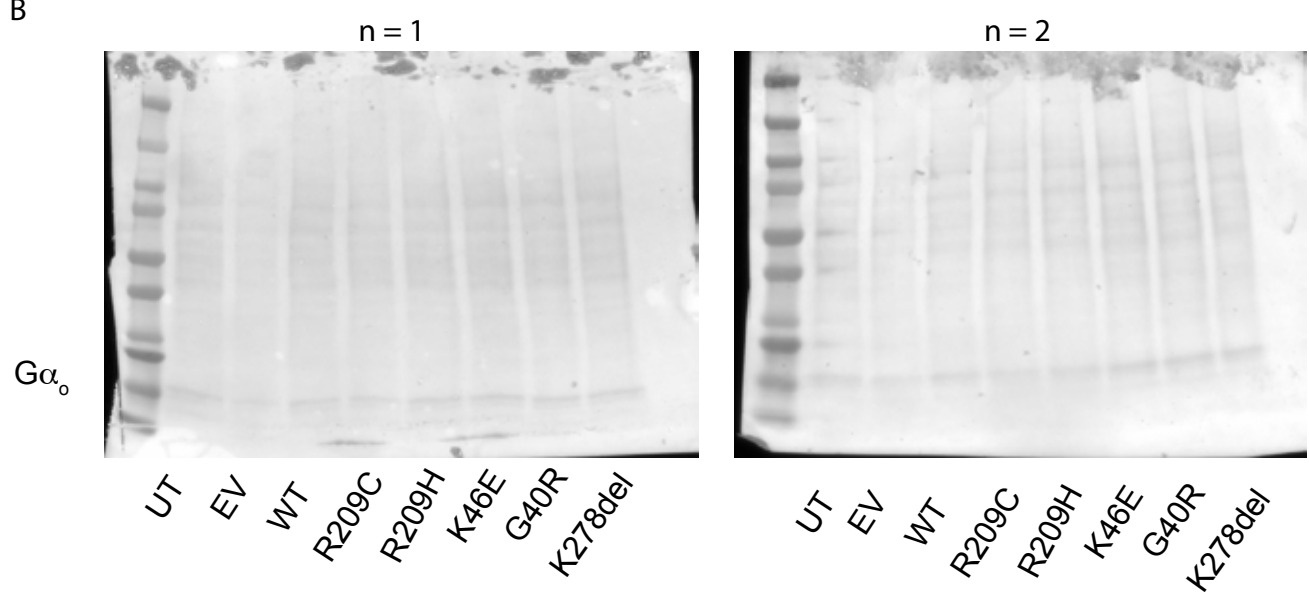

C

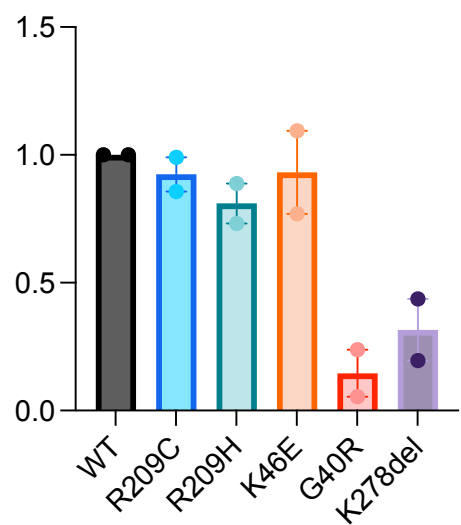

Figure S1

A

A1R

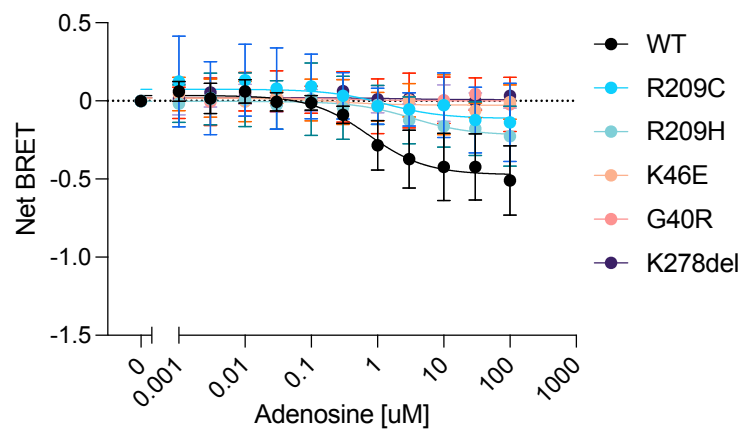

C

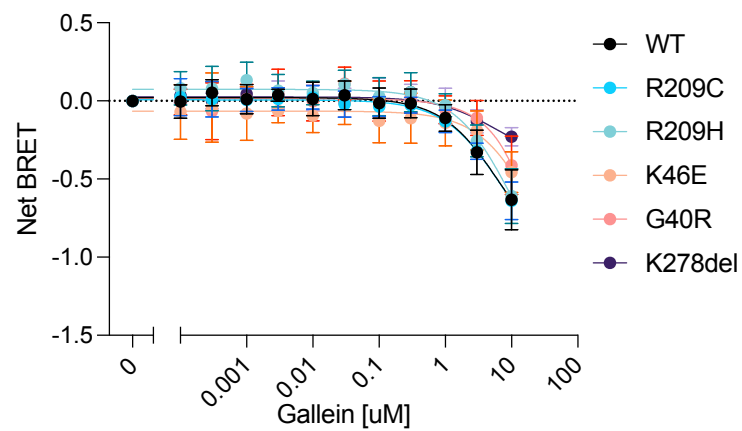

B

D2R

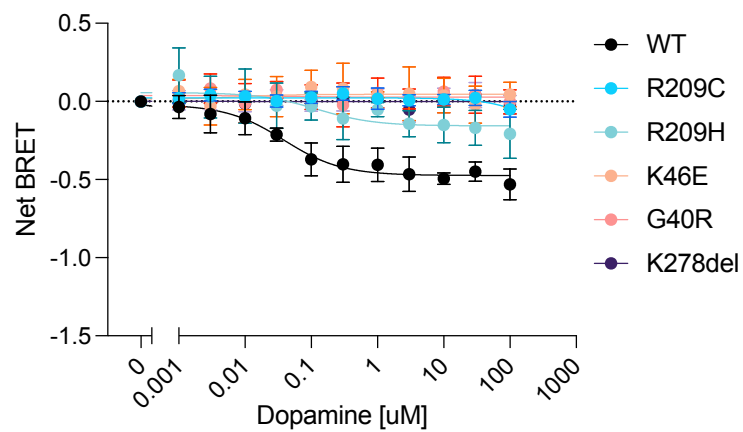

D

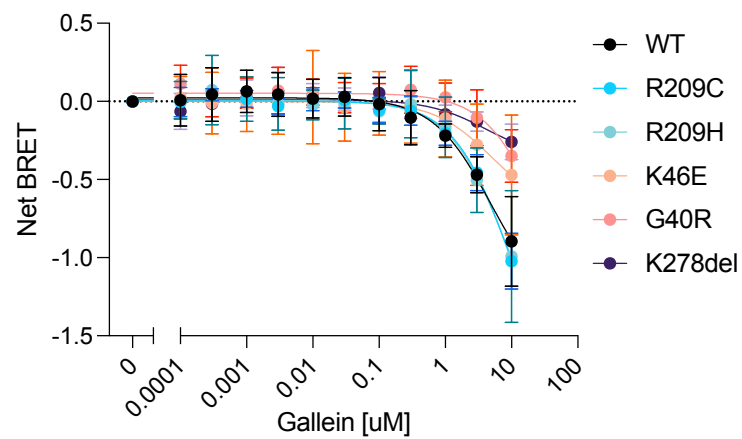

Figure S2

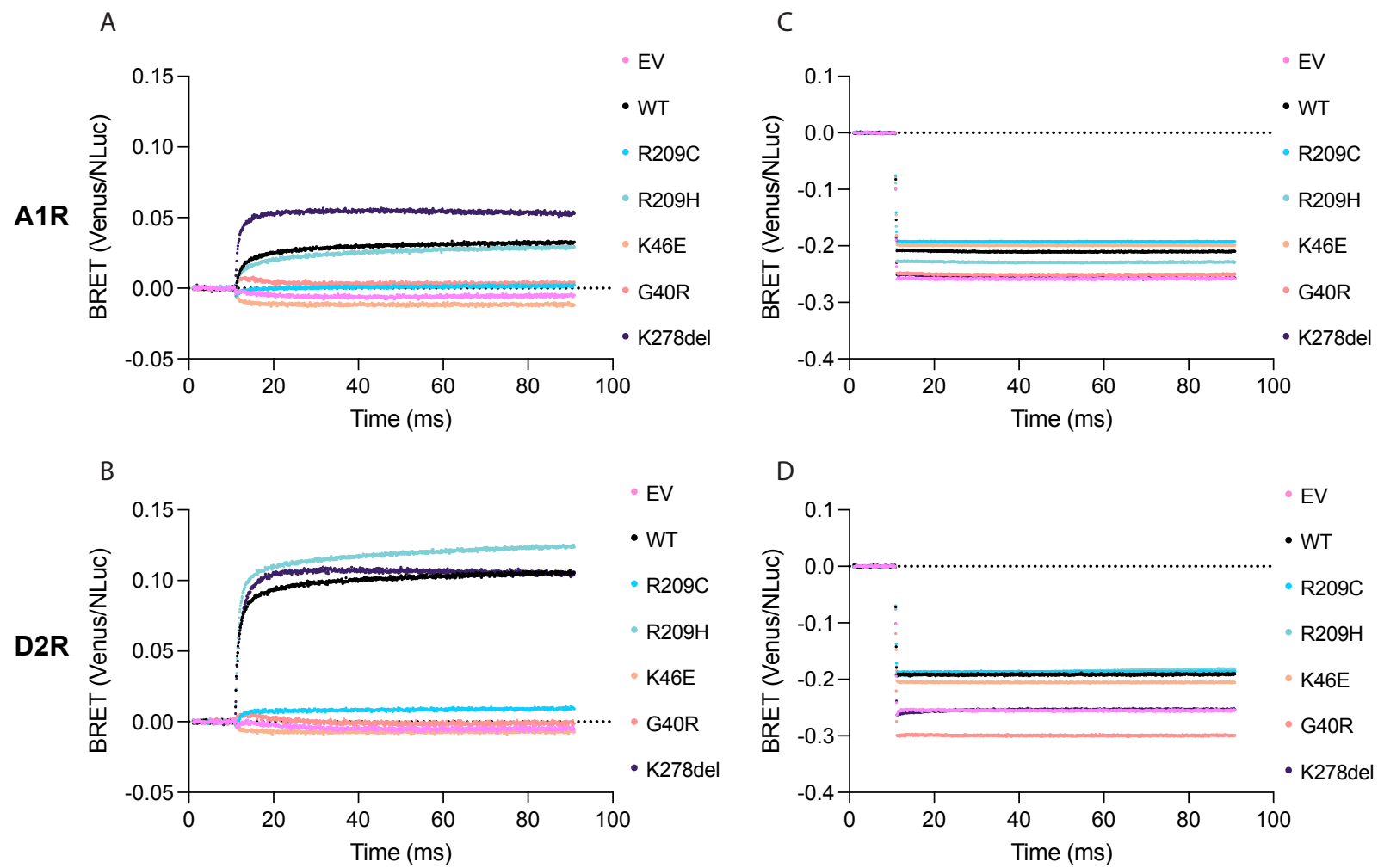

Figure S3

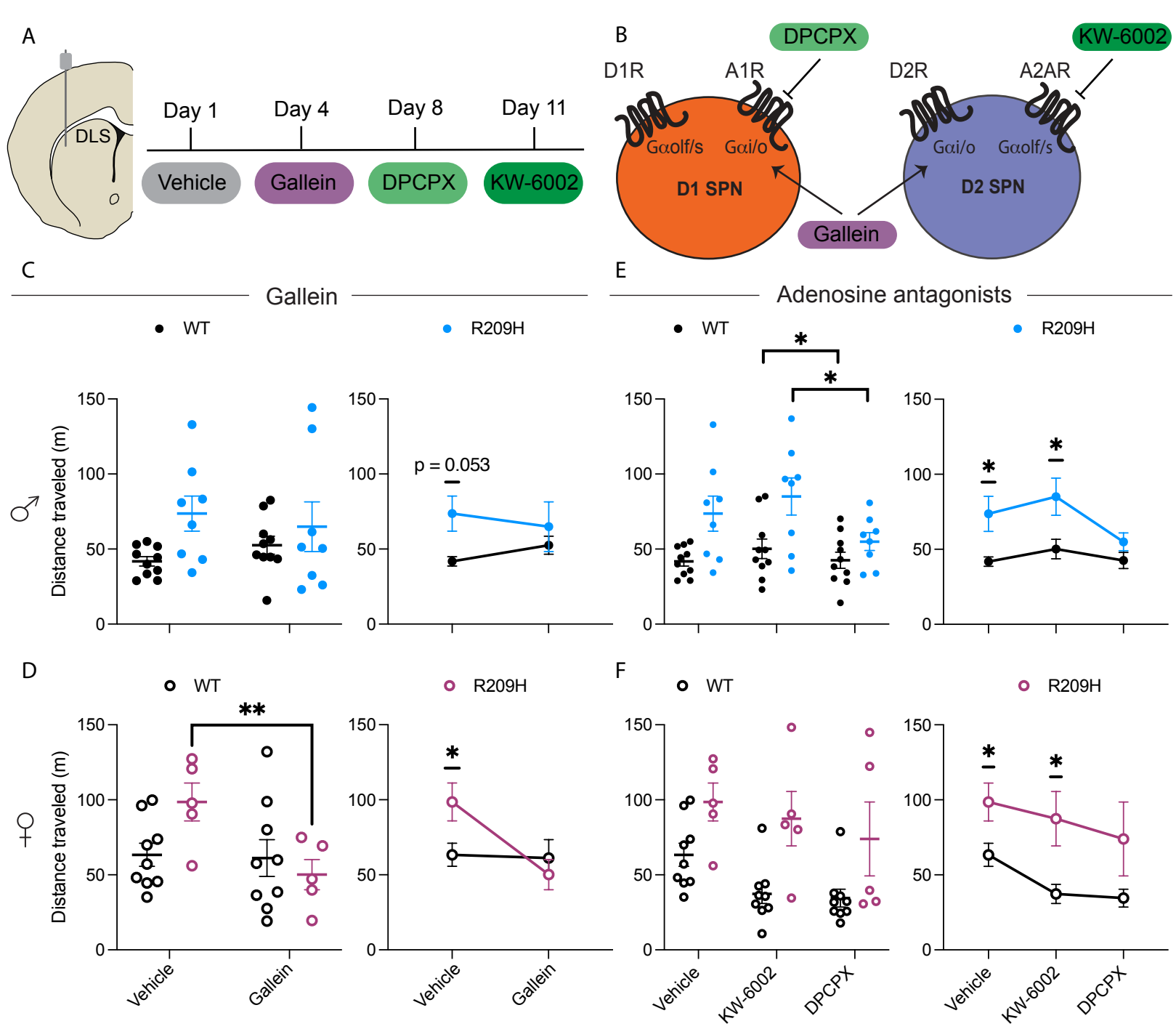

Figure S4

**Figure S1. Expression levels of WT and mutated $G\alpha_o$.** (A) Western blot from two biological replicates. (B) Ponceau stain from the two biological replicates. (C) Quantification of band intensities from Western blots. From left to right: untransfected control (UT), empty vector (EV), wild type (WT), R209C, R209H, K46E, G40R, and K278del.

**Figure S2. TRUPATH dose-response curves.** $G\alpha_o$ variants' response to (A) adenosine in cells expressing the adenosine A1 receptor or (B) dopamine in cells expressing the dopamine D2 receptor. $G\alpha_o$ variants' response to gallein was applied to cells (C) expressing the adenosine A1 receptor or (D) cells expressing the dopamine D2 receptor. K46E, G40R, and K278del were used as controls. K46E is expressed at WT levels but does not respond to agonist, and G40R and K278del are positive controls for poorly expressed $G\alpha_o$ variants (Fig. S1). Bars represent mean ± SD with individual replicates shown. All experiments have at least 3 biological replicates with 2 technical replicates.

**Figure S3. $G\beta\gamma$-GRK BRET curves.** $G\alpha_o$ variants' paired with the (A) adenosine A1 receptor in response to adenosine or (B) dopamine D2 receptor in response to dopamine. $G\alpha_o$ variants' response to gallein was applied to cells (C) expressing the adenosine A1 receptor or (D) cells expressing the dopamine D2 receptor. K46E, G40R, and K278del were used as controls. K46E is expressed at WT levels but does not respond to agonist, and G40R and K278del are positive controls for poorly expressed $G\alpha_o$ variants (Fig. S1). Bars represent mean ± SD with individual replicates shown. All experiments have at least 3 biological replicates with 3 technical replicates except for G40R, which had one technical replicate excluded due to an overgrown well.

**Figure S4. Intra-DLS infusion of gallein and selective adenosine receptor antagonists reduces hyperlocomotion in $GNAO1^{R209H/+}$ mice.** (A) Left: schematic of bilateral cannula placement targeting the dorsolateral striatum (DLS). Right: experimental timeline for the cannula infusion regimen. The same cohort of cannulated animals received all four treatments sequentially. Data in panels C and D are derived from the same animals. (B) Schematic of adenosine and dopamine receptor–G protein coupling on D1 SPNs (orange) and D2 SPNs (blue), illustrating the targets of gallein ($G\beta\gamma$ inhibitor), DPCPX (A1R antagonist), and KW-6002 (A2AR antagonist). Effect of intra-DLS gallein infusion on locomotor activity in (C) male WT and $GNAO1^{R209H/+}$ mice (WT, $n = 10$; R209H, $n = 8$) and (D) female WT and $GNAO1^{R209H/+}$ mice (WT, $n = 9$; R209H, $n = 5$). Left panels show individual data points; right panels show paired means ± SEM. are shown in the top panels. Effect of intra-DLS KW-6002 and DPCPX infusion on locomotor activity in (E) male WT and $GNAO1^{R209H/+}$ mice (WT, $n = 10$; R209H, $n = 8$) and (F) female WT and $GNAO1^{R209H/+}$ mice (WT, $n = 9$; R209H, $n = 5$). Left panels show individual data points; right panels show paired means ± SEM. Two-way repeated-measures ANOVA with Tukey's post hoc test. F statistics and post hoc test p-values can be found in Tables S10 and S11, respectively. \*$p < 0.05$, \*\*$p < 0.01$, \*\*\*$p < 0.001$.

Supplemental references:

(1) Wirth, T.; Garone, G.; Kurian, M. A.; Piton, A.; Millan, F.; Telegrafi, A.; Drouot, N.; Rudolf, G.; Chelly, J.; Marks, W.; Burglen, L.; Demailly, D.; Coubes, P.; Castro-Jimenez, M.; Joriot, S.;

- Ghoumid, J.; Belin, J.; Faucheux, J.-M.; Blumkin, L.; Hull, M.; Parnes, M.; Ravelli, C.; Poulen, G.; Calmels, N.; Nemeth, A. H.; Smith, M.; Barnicoat, A.; Ewencyk, C.; Méneret, A.; Roze, E.; Keren, B.; Mignot, C.; Beroud, C.; Jr, F. A.; Nowak, C.; Wilson, W. G.; Steel, D.; Capuano, A.; Vidailhet, M.; Lin, J.-P.; Tranchant, C.; Cif, L.; Doummar, D.; Anheim, M. Highlighting the Dystonic Phenotype Related to GNAO1. *Movement Disorders* **2022**, *37* (7), 1547–1554. https://doi.org/10.1002/mds.29074.
(2) McRae, J. F.; Clayton, S.; Fitzgerald, T. W.; Kaplanis, J.; Prigmore, E.; Rajan, D.; Sifrim, A.; Aitken, S.; Akawi, N.; Alvi, M.; Ambridge, K.; Barrett, D. M.; Bayzatinova, T.; Jones, P.; Jones, W. D.; King, D.; Krishnappa, N.; Mason, L. E.; Singh, T.; Tivey, A. R.; Ahmed, M.; Anjum, U.; Archer, H.; Armstrong, R.; Awada, J.; Balasubramanian, M.; Banka, S.; Baralle, D.; Barnicoat, A.; Batstone, P.; Baty, D.; Bennett, C.; Berg, J.; Bernhard, B.; Bevan, A. P.; Bitner-Glindzicz, M.; Blair, E.; Blyth, M.; Bohanna, D.; Bourdon, L.; Bourn, D.; Bradley, L.; Brady, A.; Brent, S.; Brewer, C.; Brunstrom, K.; Bunyan, D. J.; Burn, J.; Canham, N.; Castle, B.; Chandler, K.; Chatzimichali, E.; Cilliers, D.; Clarke, A.; Clasper, S.; Clayton-Smith, J.; Clowes, V.; Coates, A.; Cole, T.; Colgiu, I.; Collins, A.; Collinson, M. N.; Connell, F.; Cooper, N.; Cox, H.; Cresswell, L.; Cross, G.; Crow, Y.; D'Alessandro, M.; Dabir, T.; Davidson, R.; Davies, S.; de Vries, D.; Dean, J.; Deshpande, C.; Devlin, G.; Dixit, A.; Dobbie, A.; Donaldson, A.; Donnai, D.; Donnelly, D.; Donnelly, C.; Douglas, A.; Douzgou, S.; Duncan, A.; Eason, J.; Ellard, S.; Ellis, I.; Elmslie, F.; Evans, K.; Everest, S.; Fendick, T.; Fisher, R.; Flinter, F.; Foulds, N.; Fry, A.; Fryer, A.; Gardiner, C.; Gaunt, L.; Ghali, N.; Gibbons, R.; Gill, H.; Goodship, J.; Goudie, D.; Gray, E.; Green, A.; Greene, P.; Greenhalgh, L.; Gribble, S.; Harrison, R.; Harrison, L.; Harrison, V.; Hawkins, R.; He, L.; Hellens, S.; Henderson, A.; Hewitt, S.; Hildyard, L.; Hobson, E.; Holden, S.; Holder, M.; Holder, S.; Hollingsworth, G.; Homfray, T.; Humphreys, M.; Hurst, J.; Hutton, B.; Ingram, S.; Irving, M.; Islam, L.; Jackson, A.; Jarvis, J.; Jenkins, L.; Johnson, D.; Jones, E.; Josifova, D.; Joss, S.; Kaemba, B.; Kazembe, S.; Kelsell, R.; Kerr, B.; Kingston, H.; Kini, U.; Kinning, E.; Kirby, G.; Kirk, C.; Kivuva, E.; Kraus, A.; Kumar, D.; Kumar, V. K. A.; Lachlan, K.; Lam, W.; Lampe, A.; Langman, C.; Lees, M.; Lim, D.; Longman, C.; Lowther, G.; Lynch, S. A.; Magee, A.; Maher, E.; Male, A.; Mansour, S.; Marks, K.; Martin, K.; Maye, U.; McCann, E.; McConnell, V.; McEntagart, M.; McGowan, R.; McKay, K.; McKee, S.; McMullan, D. J.; McNerlan, S.; McWilliam, C.; Mehta, S.; Metcalfe, K.; Middleton, A.; Miedzybrodzka, Z.; Miles, E.; Mohammed, S.; Montgomery, T.; Moore, D.; Morgan, S.; Morton, J.; Mugalaasi, H.; Murday, V.; Murphy, H.; Naik, S.; Nemeth, A.; Nevitt, L.; Newbury-Ecob, R.; Norman, A.; O'Shea, R.; Ogilvie, C.; Ong, K.-R.; Park, S.-M.; Parker, M. J.; Patel, C.; Paterson, J.; Payne, S.; Perrett, D.; Phipps, J.; Pilz, D. T.; Pollard, M.; Pottinger, C.; Poulton, J.; Pratt, N.; Prescott, K.; Price, S.; Pridham, A.; Procter, A.; Purnell, H.; Quarrell, O.; Ragge, N.; Rahbari, R.; Randall, J.; Rankin, J.; Raymond, L.; Rice, D.; Robert, L.; Roberts, E.; Roberts, J.; Roberts, P.; Roberts, G.; Ross, A.; Rosser, E.; Saggat, A.; Samant, S.; Sampson, J.; Sandford, R.; Sarkar, A.; Schweiger, S.; Scott, R.; Scurr, I.; Selby, A.; Seller, A.; Sequeira, C.; Shannon, N.; Sharif, S.; Shaw-Smith, C.; Shearing, E.; Shears, D.; Sheridan, E.; Simonin, I.; Singzon, R.; Skitt, Z.; Smith, A.; Smith, K.; Smithson, S.; Sneddon, L.; Splitt, M.; Squires, M.; Stewart, F.; Stewart, H.; Straub, V.; Suri, M.; Sutton, V.; Swaminathan, G. J.; Sweeney, E.; Tatton-Brown, K.; Taylor, C.; Taylor, R.; Tein, M.; Temple, I. K.; Thomson, J.; Tischkowitz, M.; Tomkins, S.; Torokwa, A.; Treacy, B.; Turner, C.; Turnpenny, P.; Tysoe, C.; Vandersteen, A.; Varghese, V.; Vasudevan, P.; Vijayarangakannan, P.; Vogt, J.; Wakeling, E.; Wallwark, S.; Waters, J.; Weber, A.; Wellesley, D.; Whiteford, M.; Widaa, S.; Wilcox, S.; Wilkinson, E.; Williams, D.; Williams, N.; Wilson, L.; Woods, G.; Wragg, C.; Wright, M.; Yates, L.; Yau, M.; Nellaker, C.; Parker, M.; Firth, H. V.; Wright, C. F.; Deciphering Developmental Disorders Study. Prevalence and Architecture of de Novo Mutations in Developmental Disorders. *Nature* **2017**, *542* (7642), 433–438. https://doi.org/10.1038/nature21062.
(3) Bruun, T. U. J.; DesRoches, C.-L.; Wilson, D.; Chau, V.; Nakagawa, T.; Yamasaki, M.; Hasegawa, S.; Fukao, T.; Marshall, C.; Mercimek-Andrews, S. Prospective Cohort Study for

Identification of Underlying Genetic Causes in Neonatal Encephalopathy Using Whole-Exome Sequencing. *Genetics in Medicine* **2018**, 20 (5), 486–494. https://doi.org/10.1038/gim.2017.129.

(4) Fitzgerald, T. W.; Gerety, S. S.; Jones, W. D.; van Kogelenberg, M.; King, D. A.; McRae, J.; Morley, K. I.; Parthiban, V.; Al-Turki, S.; Ambridge, K.; Barrett, D. M.; Bayzetinova, T.; Clayton, S.; Coomber, E. L.; Gribble, S.; Jones, P.; Krishnappa, N.; Mason, L. E.; Middleton, A.; Miller, R.; Prigmore, E.; Rajan, D.; Sifrim, A.; Tivey, A. R.; Ahmed, M.; Akawi, N.; Andrews, R.; Anjum, U.; Archer, H.; Armstrong, R.; Balasubramanian, M.; Banerjee, R.; Baralle, D.; Batstone, P.; Baty, D.; Bennett, C.; Berg, J.; Bernhard, B.; Bevan, A. P.; Blair, E.; Blyth, M.; Bohanna, D.; Bourdon, L.; Bourn, D.; Brady, A.; Bragin, E.; Brewer, C.; Brueton, L.; Brunstrom, K.; Bumpstead, S. J.; Bunyan, D. J.; Burn, J.; Burton, J.; Canham, N.; Castle, B.; Chandler, K.; Clasper, S.; Clayton-Smith, J.; Cole, T.; Collins, A.; Collinson, M. N.; Connell, F.; Cooper, N.; Cox, H.; Cresswell, L.; Cross, G.; Crow, Y.; D'Alessandro, M.; Dabir, T.; Davidson, R.; Davies, S.; Dean, J.; Deshpande, C.; Devlin, G.; Dixit, A.; Dominiczak, A.; Donnelly, C.; Donnelly, D.; Douglas, A.; Duncan, A.; Eason, J.; Edkins, S.; Ellard, S.; Ellis, P.; Elmslie, F.; Evans, K.; Everest, S.; Fendick, T.; Fisher, R.; Flinter, F.; Foulds, N.; Fryer, A.; Fu, B.; Gardiner, C.; Gaunt, L.; Ghali, N.; Gibbons, R.; Gomes Pereira, S. L.; Goodship, J.; Goudie, D.; Gray, E.; Greene, P.; Greenhalgh, L.; Harrison, L.; Hawkins, R.; Hellens, S.; Henderson, A.; Hobson, E.; Holden, S.; Holder, S.; Hollingsworth, G.; Homfray, T.; Humphreys, M.; Hurst, J.; Ingram, S.; Irving, M.; Jarvis, J.; Jenkins, L.; Johnson, D.; Jones, D.; Jones, E.; Josifova, D.; Joss, S.; Kaemba, B.; Kazembe, S.; Kerr, B.; Kini, U.; Kinning, E.; Kirby, G.; Kirk, C.; Kivuva, E.; Kraus, A.; Kumar, D.; Lachlan, K.; Lam, W.; Lampe, A.; Langman, C.; Lees, M.; Lim, D.; Lowther, G.; Lynch, S. A.; Magee, A.; Maher, E.; Mansour, S.; Marks, K.; Martin, K.; Maye, U.; McCann, E.; McConnell, V.; McEntagart, M.; McGowan, R.; McKay, K.; McKee, S.; McMullan, D. J.; McNerlan, S.; Mehta, S.; Metcalfe, K.; Miles, E.; Mohammed, S.; Montgomery, T.; Moore, D.; Morgan, S.; Morris, A.; Morton, J.; Mugalaasi, H.; Murday, V.; Nevitt, L.; Newbury-Ecob, R.; Norman, A.; O'Shea, R.; Ogilvie, C.; Park, S.; Parker, M. J.; Patel, C.; Paterson, J.; Payne, S.; Phipps, J.; Pilz, D. T.; Porteous, D.; Pratt, N.; Prescott, K.; Price, S.; Pridham, A.; Procter, A.; Purnell, H.; Ragge, N.; Rankin, J.; Raymond, L.; Rice, D.; Robert, L.; Roberts, E.; Roberts, G.; Roberts, J.; Roberts, P.; Ross, A.; Rosser, E.; Saggat, A.; Samant, S.; Sandford, R.; Sarkar, A.; Schweiger, S.; Scott, C.; Scott, R.; Selby, A.; Seller, A.; Sequeira, C.; Shannon, N.; Sharif, S.; Shaw-Smith, C.; Shearing, E.; Shears, D.; Simonic, I.; Simpkin, D.; Singzon, R.; Skitt, Z.; Smith, A.; Smith, B.; Smith, K.; Smithson, S.; Sneddon, L.; Splitt, M.; Squires, M.; Stewart, F.; Stewart, H.; Suri, M.; Sutton, V.; Swaminathan, G. J.; Sweeney, E.; Tatton-Brown, K.; Taylor, C.; Taylor, R.; Tein, M.; Temple, I. K.; Thomson, J.; Tolmie, J.; Torokwa, A.; Treacy, B.; Turner, C.; Turnpenny, P.; Tysoe, C.; Vandersteen, A.; Vasudevan, P.; Vogt, J.; Wakeling, E.; Walker, D.; Waters, J.; Weber, A.; Wellesley, D.; Whiteford, M.; Widaa, S.; Wilcox, S.; Williams, D.; Williams, N.; Woods, G.; Wragg, C.; Wright, M.; Yang, F.; Yau, M.; Carter, N. P.; Parker, M.; Firth, H. V.; FitzPatrick, D. R.; Wright, C. F.; Barrett, J. C.; Hurles, M. E.; The Deciphering Developmental Disorders Study. Large-Scale Discovery of Novel Genetic Causes of Developmental Disorders. *Nature* **2015**, 519 (7542), 223–228. https://doi.org/10.1038/nature14135.
(5) Lecoquierre, F.; Duffourd, Y.; Vitobello, A.; Bruel, A.-L.; Urteaga, B.; Coubes, C.; Garret, P.; Nambot, S.; Chevarin, M.; Jouan, T.; Moutton, S.; Tran-Mau-Them, F.; Philippe, C.; Sorlin, A.; Faivre, L.; Thauvin-Robinet, C. Variant Recurrence in Neurodevelopmental Disorders: The Use of Publicly Available Genomic Data Identifies Clinically Relevant Pathogenic Missense Variants. *Genetics in Medicine* **2019**, 21 (11), 2504–2511. https://doi.org/10.1038/s41436-019-0518-x.
(6) Danti, F. R.; Galosi, S.; Romani, M.; Montomoli, M.; Carss, K. J.; Raymond, F. L.; Parrini, E.; Bianchini, C.; McShane, T.; Dale, R. C.; Mohammad, S. S.; Shah, U.; Mahant, N.; Ng, J.; McTague, A.; Samanta, R.; Vadlamani, G.; Valente, E. M.; Leuzzi, V.; Kurian, M. A.; Guerrini,

R. GNAO1 Encephalopathy: Broadening the Phenotype and Evaluating Treatment and Outcome. *Neurol Genet* **2017**, 3 (2), e143. https://doi.org/10.1212/nxg.0000000000000143.
(7) Kelly, M.; Park, M.; Mihalek, I.; Rohtus, A.; Gramm, M.; Pérez-Palma, E.; Axen, E. T.; Hung, C. Y.; Olson, H.; Swanson, L.; Anselm, I.; Briere, L. C.; High, F. A.; Sweetser, D. A.; Network, U. D.; Kayani, S.; Snyder, M.; Calvert, S.; Scheffer, I. E.; Yang, E.; Waugh, J. L.; Lal, D.; Bodamer, O.; Poduri, A. Spectrum of Neurodevelopmental Disease Associated with the GNAO1 Guanosine Triphosphate–Binding Region. *Epilepsia* **2019**, 60 (3), 406–418. https://doi.org/10.1111/epi.14653.
(8) González, M. S.; Kloosterhuis, K.; Pol, L. van de; Baas, F.; Mikkers, H. Phenotypic Diversity in GNAO1 Patients: A Comprehensive Overview of Variants and Phenotypes. *Hum. Mutat.* **2023**, 2023 (1), e6628283. https://doi.org/10.1155/2023/6628283.
(9) Takata, A.; Nakashima, M.; Saitsu, H.; Mizuguchi, T.; Mitsuhashi, S.; Takahashi, Y.; Okamoto, N.; Osaka, H.; Nakamura, K.; Tohyama, J.; Haginoya, K.; Takeshita, S.; Kuki, I.; Okanishi, T.; Goto, T.; Sasaki, M.; Sakai, Y.; Miyake, N.; Miyatake, S.; Tsuchida, N.; Iwama, K.; Minase, G.; Sekiguchi, F.; Fujita, A.; Imagawa, E.; Koshimizu, E.; Uchiyama, Y.; Hamanaka, K.; Ohba, C.; Itai, T.; Aoi, H.; Saida, K.; Sakaguchi, T.; Den, K.; Takahashi, R.; Ikeda, H.; Yamaguchi, T.; Tsukamoto, K.; Yoshitomi, S.; Oboshi, T.; Imai, K.; Kimizu, T.; Kobayashi, Y.; Kubota, M.; Kashii, H.; Baba, S.; Iai, M.; Kira, R.; Hara, M.; Ohta, M.; Miyata, Y.; Miyata, R.; Takanashi, J.; Matsui, J.; Yokochi, K.; Shimono, M.; Amamoto, M.; Takayama, R.; Hirabayashi, S.; Aiba, K.; Matsumoto, H.; Nabatame, S.; Shiihara, T.; Kato, M.; Matsumoto, N. Comprehensive Analysis of Coding Variants Highlights Genetic Complexity in Developmental and Epileptic Encephalopathy. *Nat Commun* **2019**, 10 (1), 2506. https://doi.org/10.1038/s41467-019-10482-9.
(10) Yang, X.; Niu, X.; Yang, Y.; Cheng, M.; Zhang, J.; Chen, J.; Yang, Z.; Zhang, Y. Phenotypes of GNAO1 Variants in a Chinese Cohort. *Front Neurol* **2021**, 12, 662162. https://doi.org/10.3389/fneur.2021.662162.
(11) Rim, J. H.; Kim, S. H.; Hwang, I. S.; Kwon, S. S.; Kim, J.; Kim, H. W.; Cho, M. J.; Ko, A.; Youn, S. E.; Kim, J.; Lee, Y. M.; Chung, H. J.; Lee, J. S.; Kim, H. D.; Choi, J. R.; Lee, S.-T.; Kang, H.-C. Efficient Strategy for the Molecular Diagnosis of Intractable Early-Onset Epilepsy Using Targeted Gene Sequencing. *BMC Med Genomics* **2018**, 11 (1), 6. https://doi.org/10.1186/s12920-018-0320-7.
(12) Liu, Y.; Zhang, Q.; Wang, J.; Liu, J.; Yang, W.; Yan, X.; Ouyang, Y.; Yang, H. Both Subthalamic and Pallidal Deep Brain Stimulation Are Effective for GNAO1-Associated Dystonia: Three Case Reports and a Literature Review. *Ther Adv Neurol Disord* **2022**, 15, 17562864221093507. https://doi.org/10.1177/17562864221093507.
(13) Zhu, X.; Petrovski, S.; Xie, P.; Ruzzo, E. K.; Lu, Y.-F.; McSweeney, K. M.; Ben-Zeev, B.; Nissenkorn, A.; Anikster, Y.; Oz-Levi, D.; Dhindsa, R. S.; Hitomi, Y.; Schoch, K.; Spillmann, R. C.; Heimer, G.; Marek-Yagel, D.; Tzadok, M.; Han, Y.; Worley, G.; Goldstein, J.; Jiang, Y.-H.; Lancet, D.; Pras, E.; Shashi, V.; McHale, D.; Need, A. C.; Goldstein, D. B. Whole-Exome Sequencing in Undiagnosed Genetic Diseases: Interpreting 119 Trios. *Genet Med* **2015**, 17 (10), 774–781. https://doi.org/10.1038/gim.2014.191.
(14) Powis, Z.; Towne, M. C.; Hagman, K. D. F.; Blanco, K.; Palmaer, E.; Castro, A.; Sajan, S. A.; Radtke, K.; Feyma, T. J.; Juliette, K.; Tang, S.; Sidiropoulos, C. Clinical Diagnostic Exome Sequencing in Dystonia: Genetic Testing Challenges for Complex Conditions. *Clinical Genetics* **2020**, 97 (2), 305–311. https://doi.org/10.1111/cge.13657.
(15) Gawlinski, P.; Posmyk, R.; Gambin, T.; Sielicka, D.; Chorazy, M.; Nowakowska, B.; Jhangiani, S. N.; Muzny, D. M.; Bekiesinska-Figatowska, M.; Bal, J.; Boerwinkle, E.; Gibbs, R. A.; Lupski, J. R.; Wiszniewski, W. PEHO Syndrome May Represent Phenotypic Expansion at the Severe End of the Early-Onset Encephalopathies. *Pediatr Neurol* **2016**, 60, 83–87. https://doi.org/10.1016/j.pediatrneurol.2016.03.011.

(16) Thiel, M.; Bamborschke, D.; Janzarik, W. G.; Assmann, B.; Zittel, S.; Patzer, S.; Auhuber, A.; Opp, J.; Matzker, E.; Bevot, A.; Seeger, J.; Baalen, A. van; Stüve, B.; Brockmann, K.; Cirak, S.; Koy, A. Genotype–Phenotype Correlation and Treatment Effects in Young Patients with GNAO1-Associated Disorders. *J Neurol Neurosurg Psychiatry* **2023**, *94* (10), 806–815. https://doi.org/10.1136/jnnp-2022-330261.
(17) Solis, G. P.; Kozhanova, T. V.; Koval, A.; Zhilina, S. S.; Mescheryakova, T. I.; Abramov, A. A.; Ishmuratov, E. V.; Bolshakova, E. S.; Osipova, K. V.; Ayvazyan, S. O.; Lebon, S.; Kanivets, I. V.; Pyankov, D. V.; Troccaz, S.; Silachev, D. N.; Zavadenko, N. N.; Prityko, A. G.; Katanaev, V. L. Pediatric Encephalopathy: Clinical, Biochemical and Cellular Insights into the Role of Gln52 of GNAO1 and GNAI1 for the Dominant Disease. *Cells* **2021**, *10* (10), 2749. https://doi.org/10.3390/cells10102749.
(18) Helbig, I.; Barcia, G.; Pendziwiat, M.; Ganesan, S.; Mueller, S. H.; Helbig, K. L.; Vaidiswaran, P.; Xian, J.; Galer, P. D.; Afawi, Z.; Specchio, N.; Kluger, G.; Kühlenbäumer, G.; Appenzeller, S.; Wittig, M.; Kramer, U.; van Baalen, A.; Nabbout, R. Whole-exome and HLA Sequencing in Febrile Infection-related Epilepsy Syndrome. *Ann Clin Transl Neurol* **2020**, *7* (8), 1429–1435. https://doi.org/10.1002/acn3.51062.
(19) Bobylova, M. Y.; Volkov, I. V.; Gumennik, E. V.; Rachmanina, O. A.; Abramov, M. O.; Volkova, O. K.; Bayborina, T. S.; Petrukhin, A. S. [Encephalopathy GNAO1]. *Zh Nevrol Psikhiatr Im S S Korsakova* **2023**, *123* (1), 122–130. https://doi.org/10.17116/jnevro2023123011122.
(20) Nakamura, K.; Kodera, H.; Akita, T.; Shiina, M.; Kato, M.; Hoshino, H.; Terashima, H.; Osaka, H.; Nakamura, S.; Tohyama, J.; Kumada, T.; Furukawa, T.; Iwata, S.; Shiihara, T.; Kubota, M.; Miyatake, S.; Koshimizu, E.; Nishiyama, K.; Nakashima, M.; Tsurusaki, Y.; Miyake, N.; Hayasaka, K.; Ogata, K.; Fukuda, A.; Matsumoto, N.; Saitsu, H. De Novo Mutations in GNAO1, Encoding a Gao Subunit of Heterotrimeric G Proteins, Cause Epileptic Encephalopathy. *The American Journal of Human Genetics* **2013**, *93* (3), 496–505. https://doi.org/10.1016/j.ajhg.2013.07.014.
(21) Shah, S. Phenotypic Spectrum of GNAO1 Mutation, 2022.
(22) Muir, A. M.; Myers, C. T.; Nguyen, N. T.; Saykally, J.; Craiu, D.; De Jonghe, P.; Helbig, I.; Hoffman-Zacharska, D.; Guerrini, R.; Lehesjoki, A.-E.; Marini, C.; Møller, R. S.; Serratosa, J.; Štěrbová, K.; Striano, P.; von Spiczak, S.; Weckhuysen, S.; Mefford, H. C. Genetic Heterogeneity in Infantile Spasms. *Epilepsy Research* **2019**, *156*, 106181. https://doi.org/10.1016/j.eplepsyres.2019.106181.
(23) Graziola, F.; Garone, G.; Stregapede, F.; Bosco, L.; Vigeveno, F.; Curatolo, P.; Bertini, E.; Travaglini, L.; Capuano, A. Diagnostic Yield of a Targeted Next-Generation Sequencing Gene Panel for Pediatric-Onset Movement Disorders: A 3-Year Cohort Study. *Front Genet* **2019**, *10*, 1026. https://doi.org/10.3389/fgene.2019.01026.
(24) Turro, E.; Astle, W. J.; Megy, K.; Gräf, S.; Greene, D.; Shamardina, O.; Allen, H. L.; Sanchis-Juan, A.; Frontini, M.; Thys, C.; Stephens, J.; Mapeta, R.; Burren, O. S.; Downes, K.; Haimel, M.; Tuna, S.; Aitman, T. J.; Bennett, D. L.; Calleja, P.; Carss, K.; Caulfield, M. J.; Chinnery, P. F.; Dixon, P. H.; Gale, D. P.; James, R.; Koziell, A.; Laffan, M. A.; Levine, A. P.; Maher, E. R.; Markus, H. S.; Morales, J.; Morrell, N. W.; Mumford, A. D.; Ormondroyd, E.; Rankin, S.; Rendon, A.; Richardson, S.; Roberts, I.; Roy, N. B.; Saleem, M. A.; Smith, K. G. C.; Stark, H.; Tan, R. Y. Y.; Themistocleous, A. C.; Thrasher, A. J.; Watkins, H.; Webster, A. R.; Wilkins, M. R.; Williamson, C.; Whitworth, J.; Humphray, S.; Bentley, D. R.; Kingston, N.; Walker, N.; Bradley, J. R.; Ashford, S.; Penkett, C. J.; Freson, K.; Stirrups, K. E.; Raymond, F. L.; Ouwehand, W. H. Whole-Genome Sequencing of Patients with Rare Diseases in a National Health System. *Nature* **2020**, *583* (7814), 96–102. https://doi.org/10.1038/s41586-020-2434-2.
(25) Marcé-Grau, A.; Dalton, J.; López-Pisón, J.; García-Jiménez, M. C.; Monge-Galindo, L.; Cuenca-León, E.; Giraldo, J.; Macaya, A. GNAO1 Encephalopathy: Further Delineation of a

- Severe Neurodevelopmental Syndrome Affecting Females. *Orphanet J Rare Dis* **2016**, *11*, 38. https://doi.org/10.1186/s13023-016-0416-0.
(26) Zou, D.; Wang, L.; Liao, J.; Xiao, H.; Duan, J.; Zhang, T.; Li, J.; Yin, Z.; Zhou, J.; Yan, H.; Huang, Y.; Zhan, N.; Yang, Y.; Ye, J.; Chen, F.; Zhu, S.; Wen, F.; Guo, J. Genome Sequencing of 320 Chinese Children with Epilepsy: A Clinical and Molecular Study. *Brain* **2021**, *144* (12), 3623–3634. https://doi.org/10.1093/brain/awab233.
(27) Arisaka, A.; Nakashima, M.; Kumada, S.; Inoue, K.; Nishida, H.; Mashimo, H.; Kashii, H.; Kato, M.; Maruyama, K.; Okumura, A.; Saitsu, H.; Matsumoto, N.; Fukuda, M. Association of Early-Onset Epileptic Encephalopathy with Involuntary Movements – Case Series and Literature Review. *Epilepsy & Behavior Reports* **2021**, *15*, 100417. https://doi.org/10.1016/j.ebr.2020.100417.
(28) Arya, R.; Spaeth, C.; Gilbert, D. L.; Leach, J. L.; Holland, K. D. GNAO1-Associated Epileptic Encephalopathy and Movement Disorders: C.607G>A Variant Represents a Probable Mutation Hotspot with a Distinct Phenotype. *Epileptic Disorders* **2017**, *19* (1), 67–75. https://doi.org/10.1684/epd.2017.0888.
(29) Fernández-Marmiesse, A.; Roca, I.; Díaz-Flores, F.; Cantarín, V.; Pérez-Poyato, M. S.; Fontalba, A.; Laranjeira, F.; Quintans, S.; Moldovan, O.; Felgueroso, B.; Rodríguez-Pedreira, M.; Simón, R.; Camacho, A.; Quijada, P.; Ibanez-Mico, S.; Domingno, M. R.; Benito, C.; Calvo, R.; Pérez-Cejas, A.; Carrasco, M. L.; Ramos, F.; Couce, M. L.; Ruiz-Falcó, M. L.; Gutierrez-Solana, L.; Martínez-Atienza, M. Rare Variants in 48 Genes Account for 42% of Cases of Epilepsy With or Without Neurodevelopmental Delay in 246 Pediatric Patients. *Front Neurosci* **2019**, *13*, 1135. https://doi.org/10.3389/fnins.2019.01135.
(30) Pawłowicz, M. Probable dysfunction of GABA-A receptors in patients with GNAO1-related syndromes. *Journal of Education, Health and Sport* **2021**, *11* (12), 241–253. https://doi.org/10.12775/JEHS.2021.11.12.018.
(31) Retterer, K.; Juusola, J.; Cho, M. T.; Vitazka, P.; Millan, F.; Gibellini, F.; Vertino-Bell, A.; Smaoui, N.; Neidich, J.; Monaghan, K. G.; McKnight, D.; Bai, R.; Suchy, S.; Friedman, B.; Tahiliani, J.; Pineda-Alvarez, D.; Richard, G.; Brandt, T.; Haverfield, E.; Chung, W. K.; Bale, S. Clinical Application of Whole-Exome Sequencing across Clinical Indications. *Genetics in Medicine* **2016**, *18* (7), 696–704. https://doi.org/10.1038/gim.2015.148.
(32) Saitsu, H.; Fukai, R.; Ben-Zeev, B.; Sakai, Y.; Mimaki, M.; Okamoto, N.; Suzuki, Y.; Monden, Y.; Saito, H.; Tziperman, B.; Torio, M.; Akamine, S.; Takahashi, N.; Osaka, H.; Yamagata, T.; Nakamura, K.; Tsurusaki, Y.; Nakashima, M.; Miyake, N.; Shiina, M.; Ogata, K.; Matsumoto, N. Phenotypic Spectrum of GNAO1 Variants: Epileptic Encephalopathy to Involuntary Movements with Severe Developmental Delay. *Eur J Hum Genet* **2016**, *24* (1), 129–134. https://doi.org/10.1038/ejhg.2015.92.
(33) Schirinzi, T.; Garone, G.; Travaglini, L.; Vasco, G.; Galosi, S.; Rios, L.; Castiglioni, C.; Barassi, C.; Battaglia, D.; Gambardella, M. L.; Cantonetti, L.; Graziola, F.; Marras, C. E.; Castelli, E.; Bertini, E.; Capuano, A.; Leuzzi, V. Phenomenology and Clinical Course of Movement Disorder in GNAO1 Variants: Results from an Analytical Review. *Parkinsonism & Related Disorders* **2019**, *61*, 19–25. https://doi.org/10.1016/j.parkreldis.2018.11.019.
(34) Schorling, D. C.; Dietel, T.; Evers, C.; Hinderhofer, K.; Korinthenberg, R.; Ezzo, D.; Bönnemann, C. G.; Kirschner, J. Expanding Phenotype of De Novo Mutations in GNAO1: Four New Cases and Review of Literature. *Neuropediatrics* **2017**, *48* (5), 371–377. https://doi.org/10.1055/s-0037-1603977.
(35) Xiong, J.; Peng, J.; Duan, H.-L.; Chen, C.; Wang, X.-L.; Chen, S.-M.; Yin, F. [Recurrent convulsion and pulmonary infection complicated by psychomotor retardation in an infant]. *Zhongguo Dang Dai Er Ke Za Zhi* **2018**, *20* (2), 154–157. https://doi.org/10.7499/j.issn.1008-8830.2018.02.014.
(36) Novelli, M.; Galosi, S.; Zorzi, G.; Martinelli, S.; Capuano, A.; Nardecchia, F.; Granata, T.; Pollini, L.; Rocco, M. D.; Marras, C. E.; Nardocci, N.; Leuzzi, V. GNAO1-Related Movement

(37) Koy, A.; Cirak, S.; Gonzalez, V.; Becker, K.; Roujeau, T.; Milesi, C.; Baleine, J.; Cambonie, G.; Boularan, A.; Greco, F.; Perrigault, P.-F.; Cances, C.; Dorison, N.; Doummar, D.; Roubertie, A.; Beroud, C.; Körber, F.; Stüve, B.; Waltz, S.; Mignot, C.; Nava, C.; Maarouf, M.; Coubes, P.; Cif, L. Deep Brain Stimulation Is Effective in Pediatric Patients with GNAO1 Associated Severe Hyperkinesia. *Journal of the Neurological Sciences* **2018**, 391, 31–39. https://doi.org/10.1016/j.jns.2018.05.018.
(38) Yamamoto, E. A.; Berry, M.; Harris, W.; Shahin, M. N.; Wilson, J. L.; Safarpour, D.; Raslan, A. M. Good Response to Deep Brain Stimulation in Two Forms of Inherited Chorea Related to GNAO1 and Neuroacanthocystosis with Illustrative Videos. *Mov Disord Clin Pract* **2021**, 9 (3), 401–403. https://doi.org/10.1002/mdc3.13383.
(39) Froukh, T.; Nafie, O.; Al Hait, S. A. S.; Laugwitz, L.; Sommerfeld, J.; Sturm, M.; Baraghiti, A.; Issa, T.; Al-Nazer, A.; Koch, P. A.; Hanselmann, J.; Kootz, B.; Bauer, P.; Al-Ameri, W.; Abou Jamra, R.; Alfrook, A. J.; Hamadallah, M.; Sofan, L.; Riess, A.; Haack, T. B.; Riess, O.; Buchert, R. Genetic Basis of Neurodevelopmental Disorders in 103 Jordanian Families. *Clinical Genetics* **2020**, 97 (4), 621–627. https://doi.org/10.1111/cge.13720.
(40) Kim, S. Y.; Shim, Y.; Ko, Y. J.; Park, S.; Jang, S. S.; Lim, B. C.; Kim, K. J.; Chae, J.-H. Spectrum of Movement Disorders in GNAO1 Encephalopathy: In-Depth Phenotyping and Case-by-Case Analysis. *Orphanet Journal of Rare Diseases* **2020**, 15 (1), 343. https://doi.org/10.1186/s13023-020-01594-3.
(41) Yamashita, Y.; Ogawa, T.; Ogaki, K.; Kamo, H.; Sukigara, T.; Kitahara, E.; Izawa, N.; Iwamuro, H.; Oyama, G.; Kamagata, K.; Hatano, T.; Umemura, A.; Kosaki, R.; Kubota, M.; Shimo, Y.; Hattori, N. Neuroimaging Evaluation and Successful Treatment by Using Directional Deep Brain Stimulation and Levodopa in a Patient with GNAO1-Associated Movement Disorder: A Case Report. *Journal of the Neurological Sciences* **2020**, 411, 116710. https://doi.org/10.1016/j.jns.2020.116710.
(42) Ananth, A. L.; Robichaux-Viehoever, A.; Kim, Y.-M.; Hanson-Kahn, A.; Cox, R.; Enns, G. M.; Strober, J.; Willing, M.; Schlaggar, B. L.; Wu, Y. W.; Bernstein, J. A. Clinical Course of Six Children With GNAO1 Mutations Causing a Severe and Distinctive Movement Disorder. *Pediatr. Neurol.* **2016**, 59, 81–84. https://doi.org/10.1016/j.pediatrneurol.2016.02.018.
(43) Akasaka, M.; Kamei, A.; Tanifuji, S.; Asami, M.; Ito, J.; Mizuma, K.; Oyama, K.; Tokutomi, T.; Yamamoto, K.; Fukushima, A.; Takenouchi, T.; Uehara, T.; Suzuki, H.; Kosaki, K. GNAO1 Mutation-Related Severe Involuntary Movements Treated with Gabapentin. *Brain Dev* **2021**, 43 (4), 576–579. https://doi.org/10.1016/j.braindev.2020.12.002.
(44) Brunet, T.; Jech, R.; Brugger, M.; Kovacs, R.; Alhaddad, B.; Leszinski, G.; Riedhammer, K. M.; Westphal, D. S.; Mahle, I.; Mayerhanser, K.; Skovranek, M.; Weber, S.; Graf, E.; Berutti, R.; Necpál, J.; Havránková, P.; Pavelekova, P.; Hempel, M.; Kotzaeridou, U.; Hoffmann, G. F.; Leiz, S.; Makowski, C.; Roser, T.; Schroeder, S. A.; Steinfeld, R.; Strobl-Wildemann, G.; Hoefele, J.; Borggraefe, I.; Distelmaier, F.; Strom, T. M.; Winkelmann, J.; Meitinger, T.; Zech, M.; Wagner, M. De Novo Variants in Neurodevelopmental Disorders—Experiences from a Tertiary Care Center. *Clinical Genetics* **2021**, 100 (1), 14–28. https://doi.org/10.1111/cge.13946.
(45) Danhofer, P.; Zech, M.; Bálintová, Z.; Baláž, M.; Jech, R.; Ošlejšková, H. Brittle Biballism-Dystonia in a Pediatric Patient with GNAO1 Mutation Managed Using Pallidal Deep Brain Stimulation. *Mov Disord Clin Pract* **2021**, 8 (1), 153–155. https://doi.org/10.1002/mdc3.13118.
(46) Dzinovic, I.; Škorvánek, M.; Necpál, J.; Boesch, S.; Švantnerová, J.; Wagner, M.; Havránková, P.; Pavelekova, P.; Haň, V.; Janzarik, W. G.; Berweck, S.; Diebold, I.; Kuster, A.; Jech, R.; Winkelmann, J.; Zech, M. Dystonia as a Prominent Presenting Feature in

Developmental and Epileptic Encephalopathies: A Case Series. *Parkinsonism & Related Disorders* **2021**, 90, 73–78. https://doi.org/10.1016/j.parkreldis.2021.08.007.

(47) Kwong, A. K.-Y.; Tsang, M. H.-Y.; Fung, J. L.-F.; Mak, C. C.-Y.; Chan, K. L.-S.; Rodenburg, R. J. T.; Lek, M.; Huang, S.; Pajusalu, S.; Yau, M.-M.; Tsoi, C.; Fung, S.; Liu, K.-T.; Ma, C.-K.; Wong, S.; Yau, E. K.-C.; Tai, S.-M.; Fung, E. L.-W.; Wu, N. S.-P.; Tsung, L.-Y.; Smeitink, J.; Chung, B. H.-Y.; Fung, C.-W. Exome Sequencing in Paediatric Patients with Movement Disorders. *Orphanet J Rare Dis* **2021**, 16 (1), 32. https://doi.org/10.1186/s13023-021-01688-6.
(48) Malaquias, M. J.; Fineza, I.; Loureiro, L.; Cardoso, L.; Alonso, I.; Magalhães, M. GNAO1 Mutation Presenting as Dyskinetic Cerebral Palsy. *Neurol Sci* **2019**, 40 (10), 2213–2216. https://doi.org/10.1007/s10072-019-03964-7.
(49) Waak, M.; Mohammad, S. S.; Coman, D.; Sinclair, K.; Copeland, L.; Silburn, P.; Coyne, T.; McGill, J.; O'Regan, M.; Selway, R.; Symonds, J.; Grattan-Smith, P.; Lin, J.-P.; Dale, R. C.; Malone, S. GNAO1-Related Movement Disorder with Life-Threatening Exacerbations: Movement Phenomenology and Response to DBS. *J Neurol Neurosurg Psychiatry* **2018**, 89 (2), 221–222. https://doi.org/10.1136/jnnp-2017-315653.
(50) Zech, M.; Jech, R.; Boesch, S.; Škorvánek, M.; Weber, S.; Wagner, M.; Zhao, C.; Jochim, A.; Necpál, J.; Dincer, Y.; Vill, K.; Distelmaier, F.; Stoklosa, M.; Krenn, M.; Grunwald, S.; Bock-Bierbaum, T.; Fečíková, A.; Havránková, P.; Roth, J.; Příhodová, I.; Adamovičová, M.; Ulmanová, O.; Bechyně, K.; Danhofer, P.; Veselý, B.; Haň, V.; Pavelekova, P.; Gdovinová, Z.; Mantel, T.; Meindl, T.; Sitzberger, A.; Schröder, S.; Blaschek, A.; Roser, T.; Bonfert, M. V.; Haberlandt, E.; Plecko, B.; Leineweber, B.; Berweck, S.; Herberhold, T.; Langguth, B.; Švantnerová, J.; Minár, M.; Ramos-Rivera, G. A.; Wojcik, M. H.; Pajusalu, S.; Öunap, K.; Schatz, U. A.; Pölsler, L.; Milenkovic, I.; Laccone, F.; Pilshofer, V.; Colombo, R.; Patzer, S.; Iuso, A.; Vera, J.; Troncoso, M.; Fang, F.; Prokisch, H.; Wilbert, F.; Eckenweiler, M.; Graf, E.; Westphal, D. S.; Riedhammer, K. M.; Brunet, T.; Alhaddad, B.; Berutti, R.; Strom, T. M.; Hecht, M.; Baumann, M.; Wolf, M.; Telegrafi, A.; Person, R. E.; Zamora, F. M.; Henderson, L. B.; Weise, D.; Musacchio, T.; Volkmann, J.; Szuto, A.; Becker, J.; Cremer, K.; Sycha, T.; Zimprich, F.; Kraus, V.; Makowski, C.; Gonzalez-Alegre, P.; Bardakjian, T. M.; Ozelius, L. J.; Vetro, A.; Guerrini, R.; Maier, E.; Borggraefe, I.; Kuster, A.; Wortmann, S. B.; Hackenberg, A.; Steinfeld, R.; Assmann, B.; Staufner, C.; Opladen, T.; Růžicka, E.; Cohn, R. D.; Dymont, D.; Chung, W. K.; Engels, H.; Ceballos-Baumann, A.; Ploski, R.; Daumke, O.; Haslinger, B.; Mall, V.; Oexle, K.; Winkelmann, J. Monogenic Variants in Dystonia: An Exome-Wide Sequencing Study. *Lancet Neurol* **2020**, 19 (11), 908–918. [https://doi.org/10.1016/S1474-4422(20)30312-4](https://doi.org/10.1016/S1474-4422%2820%2930312-4).
(51) Blumkin, L.; Lerman-Sagie, T.; Westenberger, A.; Ben-Pazi, H.; Zerem, A.; Yosovich, K.; Lev, D. Multiple Causes of Pediatric Early Onset Chorea-Clinical and Genetic Approach. *Neuropediatrics* **2018**, 49 (4), 246–255. https://doi.org/10.1055/s-0038-1645884.
(52) Dhamija, R.; Mink, J. W.; Shah, B. B.; Goodkin, H. P. GNAO1-Associated Movement Disorder. *Movement Disorders Clinical Practice* **2016**, 3 (6), 615–617. https://doi.org/10.1002/mdc3.12344.
(53) Kulkarni, N.; Tang, S.; Bhardwaj, R.; Bernes, S.; Grebe, T. A. Progressive Movement Disorder in Brothers Carrying a GNAO1 Mutation Responsive to Deep Brain Stimulation. *J Child Neurol* **2016**, 31 (2), 211–214. https://doi.org/10.1177/0883073815587945.
(54) Marecos, C.; Duarte, S.; Alonso, I.; Calado, E.; Moreira, A. [GNAO1: a new gene to consider on early-onset childhood dystonia]. *Rev Neurol* **2018**, 66 (9), 321–322.
(55) Menke, L. A.; Engelen, M.; Alders, M.; Odekerken, V. J. J.; Baas, F.; Cobben, J. M. Recurrent GNAO1 Mutations Associated With Developmental Delay and a Movement Disorder. *J Child Neurol* **2016**, 31 (14), 1598–1601. https://doi.org/10.1177/0883073816666474.

(56) 胡文静; 方红军; 唐静文; 周珍; 吴丽文. 早期发病的神经发育障碍伴非自主运动患儿1例的遗传学分析及文献复习. *中华医学遗传学杂志* **2023**, 40 (04), 385–389. https://doi.org/10.3760/cma.j.cn511374-20221206-00844.
(57) van der Ven, A. T.; Johannsen, J.; Kortüm, F.; Wagner, M.; Tsiakas, K.; Bierhals, T.; Lessel, D.; Herget, T.; Kloth, K.; Lisfeld, J.; Scholz, T.; Obi, N.; Wortmann, S.; Prokisch, H.; Kubisch, C.; Denecke, J.; Santer, R.; Hempel, M. Prevalence and Clinical Prediction of Mitochondrial Disorders in a Large Neuropediatric Cohort. *Clin Genet* **2021**, 100 (6), 766–770. https://doi.org/10.1111/cge.14061.
(58) Honey, C. M.; Malhotra, A. K.; Tarailo-Graovac, M.; van Karnebeek, C. D. M.; Horvath, G.; Sulistyanto, A. GNAO1 Mutation-Induced Pediatric Dystonic Storm Rescue With Pallidal Deep Brain Stimulation. *J Child Neurol* **2018**, 33 (6), 413–416. https://doi.org/10.1177/0883073818756134.
(59) Schöne-Bake, J.-C.; Auber, B.; Jack, T.; Hartmann, H. P 411. GNAO1-Associated Hyperkinetic-Dystonic Movement Disorder and Developmental Delay in a 14-Year-Old Girl. In *Neuropediatrics*; Georg Thieme Verlag KG, 2018; Vol. 49, p P411. https://doi.org/10.1055/s-0038-1675951.
(60) Matthews, A. M.; Blydt-Hansen, I.; Al-Jabri, B.; Andersen, J.; Tarailo-Graovac, M.; Price, M.; Selby, K.; Demos, M.; Connolly, M.; Drögemöller, B.; Shyr, C.; Mwenifumbo, J.; Elliott, A. M.; Lee, J.; Ghani, A.; Stöckler, S.; Salvarinova, R.; Vallance, H.; Sinclair, G.; Ross, C. J.; Wasserman, W. W.; McKinnon, M. L.; Horvath, G. A.; Goez, H.; van Karnebeek, C. D.; TIDE BC, United for Metabolic Diseases and the CAUSES Study. Atypical Cerebral Palsy: Genomics Analysis Enables Precision Medicine. *Genet Med* **2019**, 21 (7), 1621–1628. https://doi.org/10.1038/s41436-018-0376-y.
(61) Carecchio, M.; Invernizzi, F.; González-Latapi, P.; Panteghini, C.; Zorzi, G.; Romito, L.; Leuzzi, V.; Galosi, S.; Reale, C.; Zibordi, F.; Joseph, A. P.; Topf, M.; Piano, C.; Bentivoglio, A. R.; Girotti, F.; Morana, P.; Morana, B.; Kurian, M. A.; Garavaglia, B.; Mencacci, N. E.; Lubbe, S. J.; Nardocci, N. Frequency and Phenotypic Spectrum of KMT2B Dystonia in Childhood: A Single-Center Cohort Study. *Mov Disord* **2019**, 34 (10), 1516–1527. https://doi.org/10.1002/mds.27771.
(62) May, H. J.; Fasheun, J. A.; Bain, J. M.; Baugh, E. H.; Bier, L. E.; Revah-Politi, A.; Roye, D. P.; Goldstein, D. B.; Carmel, J. B. Cerebral Palsy Genetics: Who to Test? *Dev Med Child Neurol* **2021**, 63 (12), 1448–1455. https://doi.org/10.1111/dmcn.14948.
(63) Mahfouz, N. A.; Kizhakkedath, P.; Ibrahim, A.; El Naofal, M.; Ramaswamy, S.; Harilal, D.; Qutub, Y.; Uddin, M.; Taylor, A.; Alloub, Z.; AlBanna, A.; Abuhammour, W.; Fathalla, B.; Tayoun, A. A. Utility of Clinical Exome Sequencing in a Complex Emirati Pediatric Cohort. *Comput Struct Biotechnol J* **2020**, 18, 1020–1027. https://doi.org/10.1016/j.csbj.2020.04.013.
(64) Nashabat, M.; Al Qahtani, X. S.; Almakdub, S.; Altwaijri, W.; Ba-Armah, D. M.; Hundallah, K.; Al Hashem, A.; Al Tala, S.; Maddirevula, S.; Alkuraya, F. S.; Tabarki, B.; Alfadhel, M. The Landscape of Early Infantile Epileptic Encephalopathy in a Consanguineous Population. *Seizure* **2019**, 69, 154–172. https://doi.org/10.1016/j.seizure.2019.04.018.
(65) Talvik, I.; Møller, R. S.; Vaher, M.; Vaher, U.; Larsen, L. H.; Dahl, H. A.; Ilves, P.; Talvik, T. Clinical Phenotype of De Novo GNAO1 Mutation. *Child Neurol Open* **2015**, 2 (2), 2329048X15583717. https://doi.org/10.1177/2329048X15583717.
(66) Rosello, M.; Caro-Llopis, A.; Orellana, C.; Oltra, S.; Alemany-Albert, M.; Marco-Hernandez, A. V.; Monfort, S.; Pedrola, L.; Martinez, F.; Tomás, M. Hidden Etiology of Cerebral Palsy: Genetic and Clinical Heterogeneity and Efficient Diagnosis by next-Generation Sequencing. *Pediatr Res* **2021**, 90 (2), 284–288. https://doi.org/10.1038/s41390-020-01250-3.
(67) Yilmaz, S.; Turhan, T.; Ceylaner, S.; Gökben, S.; Tekgul, H.; Serdaroglu, G. Excellent Response to Deep Brain Stimulation in a Young Girl with GNAO1-Related Progressive

Choreoathetosis. *Childs Nerv Syst* **2016**, 32 (9), 1567–1568. https://doi.org/10.1007/s00381-016-3139-6.

(68) Córdoba, M.; Rodríguez-Quiroga, S. A.; Vega, P. A.; Salinas, V.; Perez-Maturo, J.; Amartino, H.; Vásquez-Dusefante, C.; Medina, N.; González-Morón, D.; Kauffman, M. A. Whole Exome Sequencing in Neurogenetic Odysseys: An Effective, Cost- and Time-Saving Diagnostic Approach. *PLOS ONE* **2018**, 13 (2), e0191228. https://doi.org/10.1371/journal.pone.0191228.
(69) Al Masseri, Z.; AlSayed, M. Gonadal Mosaicism in GNAO1 Causing Neurodevelopmental Disorder with Involuntary Movements; Two Additional Variants. *Mol Genet Metab Rep* **2022**, 31, 100864. https://doi.org/10.1016/j.ymgmr.2022.100864.
(70) Monies, D.; Abouelhoda, M.; Assoum, M.; Moghrabi, N.; Rafiullah, R.; Almontashiri, N.; Alowain, M.; Alzaidan, H.; Alsayed, M.; Subhani, S.; Cupler, E.; Faden, M.; Alhashem, A.; Qari, A.; Chedrawi, A.; Aldhalaan, H.; Kurdi, W.; Khan, S.; Rahbeeni, Z.; Alotaibi, M.; Goljan, E.; Elbardisy, H.; ElKalioby, M.; Shah, Z.; Alruwaili, H.; Jaafar, A.; Albar, R.; Akilan, A.; Tayeb, H.; Tahir, A.; Fawzy, M.; Nasr, M.; Makki, S.; Alfaifi, A.; Akleh, H.; Yamani, S.; Bubshait, D.; Mahnashi, M.; Basha, T.; Alsagheir, A.; Khaled, M. A.; Alsaleem, K.; Almugbel, M.; Badawi, M.; Bashiri, F.; Bohlega, S.; Sulaiman, R.; Tous, E.; Ahmed, S.; Algoufi, T.; Al-Mousa, H.; Alaki, E.; Alhumaidi, S.; Alghamdi, H.; Alghamdi, M.; Sahly, A.; Nahrir, S.; Al-Ahmari, A.; Alkuraya, H.; Almehaidib, A.; Abanemai, M.; Alsohaibaini, F.; Alsaud, B.; Arnaout, R.; Abdel-Salam, G. M. H.; Aldhekri, H.; AlKhater, S.; Alqadi, K.; Alsabban, E.; Alshareef, T.; Awartani, K.; Banjar, H.; Alsahan, N.; Abosoudah, I.; Alashwal, A.; Aldekhail, W.; Alhajjar, S.; Al-Mayouf, S.; Alsemari, A.; Alshuaibi, W.; Altala, S.; Altalhi, A.; Baz, S.; Hamad, M.; Abalkhail, T.; Alenazi, B.; Alkaff, A.; Almohareb, F.; Al Mutairi, F.; Alsaleh, M.; Alsonbul, A.; Alzelaye, S.; Bahzad, S.; Manee, A. B.; Jarrad, O.; Meriki, N.; Albeirouti, B.; Alqasmi, A.; AlBalwi, M.; Makhseed, N.; Hassan, S.; Salih, I.; Salih, M. A.; Shaheen, M.; Sermin, S.; Shahrukh, S.; Hashmi, S.; Shawli, A.; Tajuddin, A.; Tamim, A.; Alnahari, A.; Ghemlas, I.; Hussein, M.; Wali, S.; Murad, H.; Meyer, B. F.; Alkuraya, F. S. Lessons Learned from Large-Scale, First-Tier Clinical Exome Sequencing in a Highly Consanguineous Population. *Am J Hum Genet* **2019**, 105 (4), 879. https://doi.org/10.1016/j.ajhg.2019.09.019.
(71) Okumura, A.; Maruyama, K.; Shibata, M.; Kurahashi, H.; Ishii, A.; Numoto, S.; Hirose, S.; Kawai, T.; Iso, M.; Kataoka, S.; Okuno, Y.; Muramatsu, H.; Kojima, S. A Patient with a GNAO1 Mutation with Decreased Spontaneous Movements, Hypotonia, and Dystonic Features. *Brain Dev* **2018**, 40 (10), 926–930. https://doi.org/10.1016/j.braindev.2018.06.005.
(72) Fung, E. L.; Mo, C.; Fung, S. T.-H.; Chan, A. Y.; Lau, K.; Chan, E. K.; Chan, D. Y.; Zhu, X.; Chan, D. T.; Poon, W. Deep Brain Stimulation in a Young Child with GNAO1 Mutation – Feasible and Helpful. *Surg Neurol Int* **2022**, 13, 285. https://doi.org/10.25259/SNI_166_2022.
(73) Ling, W.; Huang, D.; Yang, F.; Yang, Z.; Liu, M.; Zhu, Q.; Huang, J.; Zhou, R.; Chen, X. Treating GNAO1 Mutation-Related Severe Movement Disorders with Oxcarbazepine: A Case Report. *Transl Pediatr* **2022**, 11 (9), 1577–1587. https://doi.org/10.21037/tp-22-297.
(74) Miyamoto, S.; Nakashima, M.; Fukumura, S.; Kumada, S.; Saitsu, H. An Intronic GNAO1 Variant Leading to In-Frame Insertion Cause Movement Disorder Controlled by Deep Brain Stimulation. *Neurogenetics* **2022**, 23 (2), 129–135. https://doi.org/10.1007/s10048-022-00686-5.
(75) Hamdan, F. F.; Myers, C. T.; Cossette, P.; Lemay, P.; Spiegelman, D.; Laporte, A. D.; Nassif, C.; Diallo, O.; Monlong, J.; Cadieux-Dion, M.; Dobrzeniecka, S.; Meloche, C.; Retterer, K.; Cho, M. T.; Rosenfeld, J. A.; Bi, W.; Massicotte, C.; Miguët, M.; Brunga, L.; Regan, B. M.; Mo, K.; Tam, C.; Schneider, A.; Hollingsworth, G.; FitzPatrick, D. R.; Donaldson, A.; Canham, N.; Blair, E.; Kerr, B.; Fry, A. E.; Thomas, R. H.; Shelagh, J.; Hurst, J. A.; Brittain, H.; Blyth, M.; Lebel, R. R.; Gerkes, E. H.; Davis-Keppen, L.; Stein, Q.; Chung, W. K.; Dorison, S. J.; Benke, P. J.; Fassi, E.; Corsten-Janssen, N.; Kamsteeg, E.-J.; Mau-Them, F. T.; Bruel, A.-L.; Verloes, A.; Öunap, K.; Wojcik, M. H.; Albert, D. V. F.; Venkateswaran, S.; Ware, T.; Jones, D.;

- Liu, Y.-C.; Mohammad, S. S.; Bizargity, P.; Bacino, C. A.; Leuzzi, V.; Martinelli, S.; Dallapiccola, B.; Tartaglia, M.; Blumkin, L.; Wierenga, K. J.; Purcarin, G.; O'Byrne, J. J.; Stockler, S.; Lehman, A.; Keren, B.; Nougues, M.-C.; Mignot, C.; Auvin, S.; Nava, C.; Hiatt, S. M.; Bebin, M.; Shao, Y.; Scaglia, F.; Lalani, S. R.; Frye, R. E.; Jarjour, I. T.; Jacques, S.; Boucher, R.-M.; Riou, E.; Srour, M.; Carmant, L.; Lortie, A.; Major, P.; Diadori, P.; Dubeau, F.; D'Anjou, G.; Bourque, G.; Berkovic, S. F.; Sadleir, L. G.; Campeau, P. M.; Kibar, Z.; Lafrenière, R. G.; Girard, S. L.; Mercimek-Mahmutoglu, S.; Boelman, C.; Rouleau, G. A.; Scheffer, I. E.; Mefford, H. C.; Andrade, D. M.; Rossignol, E.; Minassian, B. A.; Michaud, J. L. High Rate of Recurrent De Novo Mutations in Developmental and Epileptic Encephalopathies. *Am J Hum Genet* **2017**, *101* (5), 664–685. https://doi.org/10.1016/j.ajhg.2017.09.008.
(76) Farwell, K. D.; Shahmirzadi, L.; El-Khechen, D.; Powis, Z.; Chao, E. C.; Tippin Davis, B.; Baxter, R. M.; Zeng, W.; Mroske, C.; Parra, M. C.; Gandomi, S. K.; Lu, I.; Li, X.; Lu, H.; Lu, H.-M.; Salvador, D.; Ruble, D.; Lao, M.; Fischbach, S.; Wen, J.; Lee, S.; Elliott, A.; Dunlop, C. L. M.; Tang, S. Enhanced Utility of Family-Centered Diagnostic Exome Sequencing with Inheritance Model-Based Analysis: Results from 500 Unselected Families with Undiagnosed Genetic Conditions. *Genet Med* **2015**, *17* (7), 578–586. https://doi.org/10.1038/gim.2014.154.
(77) Takezawa, Y.; Kikuchi, A.; Haginoya, K.; Niihori, T.; Numata-Uematsu, Y.; Inui, T.; Yamamura-Suzuki, S.; Miyabayashi, T.; Anzai, M.; Suzuki-Muromoto, S.; Okubo, Y.; Endo, W.; Togashi, N.; Kobayashi, Y.; Onuma, A.; Funayama, R.; Shiota, M.; Nakayama, K.; Aoki, Y.; Kure, S. Genomic Analysis Identifies Masqueraders of Full-term Cerebral Palsy. *Ann Clin Transl Neurol* **2018**, *5* (5), 538–551. https://doi.org/10.1002/acn3.551.
(78) Benato, A.; Carecchio, M.; Burlina, A.; Paoloni, F.; Sartori, S.; Nosadini, M.; d'Avella, D.; Landi, A.; Antonini, A. Long-Term Effect of Subthalamic and Pallidal Deep Brain Stimulation for Status Dystonicus in Children with Methylmalonic Acidemia and GNAO1 Mutation. *J Neural Transm (Vienna)* **2019**, *126* (6), 739–757. https://doi.org/10.1007/s00702-019-02010-2.
(79) Liu, P.; Meng, L.; Normand, E. A.; Xia, F.; Song, X.; Ghazi, A.; Rosenfeld, J.; Magoulas, P. L.; Braxton, A.; Ward, P.; Dai, H.; Yuan, B.; Bi, W.; Xiao, R.; Wang, X.; Chiang, T.; Vetrini, F.; He, W.; Cheng, H.; Dong, J.; Gijavanekar, C.; Benke, P. J.; Bernstein, J. A.; Eble, T.; Eroglu, Y.; Erwin, D.; Escobar, L.; Gibson, J. B.; Gripp, K.; Kleppe, S.; Koenig, M. K.; Lewis, A. M.; Natowicz, M.; Mancias, P.; Minor, L.; Scaglia, F.; Schaaf, C. P.; Streff, H.; Vernon, H.; Uhles, C. L.; Zackai, E. H.; Wu, N.; Sutton, V. R.; Beaudet, A. L.; Muzny, D.; Gibbs, R. A.; Posey, J. E.; Lalani, S.; Shaw, C.; Eng, C. M.; Lupski, J. R.; Yang, Y. Reanalysis of Clinical Exome Sequencing Data. *New England Journal of Medicine* **2019**, *380* (25), 2478–2480. https://doi.org/10.1056/NEJMc1812033.
(80) De Novo Mutations in Synaptic Transmission Genes Including DNM1 Cause Epileptic Encephalopathies. *Am J Hum Genet* **2017**, *100* (1), 179. https://doi.org/10.1016/j.ajhg.2016.12.012.
(81) De Novo Mutations in Synaptic Transmission Genes Including DNM1 Cause Epileptic Encephalopathies. *Am J Hum Genet* **2014**, *95* (4), 360–370. https://doi.org/10.1016/j.ajhg.2014.08.013.
(82) Yuskaitis, C. J.; Ruzhnikov, M. R. Z.; Howell, K. B.; Allen, I. E.; Kapur, K.; Dlugos, D. J.; Scheffer, I. E.; Poduri, A.; Sherr, E. H. Infantile Spasms of Unknown Cause: Predictors of Outcome and Genotype-Phenotype Correlation. *Pediatr Neurol* **2018**, *87*, 48–56. https://doi.org/10.1016/j.pediatrneurol.2018.04.012.
(83) Baldrige, D.; Heeley, J.; Vineyard, M.; Manwaring, L.; Toler, T. L.; Fassi, E.; Fiala, E.; Brown, S.; Goss, C. W.; Willing, M.; Grange, D. K.; Kozel, B. A.; Shinawi, M. The Exome Clinic and the Role of Medical Genetics Expertise in Interpretation of Exome Sequencing Results. *Genet Med* **2017**, *19* (9), 1040–1048. https://doi.org/10.1038/gim.2016.224.

(84) De Novo Mutations in SLC1A2 and CACNA1A Are Important Causes of Epileptic Encephalopathies. *Am J Hum Genet* **2016**, 99 (2), 287–298. https://doi.org/10.1016/j.ajhg.2016.06.003.
(85) Karaca, E.; Posey, J. E.; Akdemir, Z. C.; Pehlivan, D.; Harel, T.; Jhangiani, S. N.; Bayram, Y.; Song, X.; Bahrambeigi, V.; Yuregir, O. O.; Bozdogan, S.; Yesil, G.; Isikay, S.; Muzny, D.; Gibbs, R. A.; Lupski, J. R. Phenotypic Expansion Illuminates Multilocus Pathogenic Variation. *Genet Med* **2018**, 20 (12), 1528–1537. https://doi.org/10.1038/gim.2018.33.
(86) Gerald, B.; Ramsey, K.; Belnap, N.; Szelinger, S.; Siniard, A. L.; Balak, C.; Russell, M.; Richholt, R.; De Both, M.; Claasen, A. M.; Schrauwen, I.; Huentelman, M. J.; Craig, D. W.; Rangasamy, S.; Narayanan, V. Neonatal Epileptic Encephalopathy Caused by de Novo GNAO1 Mutation Misdiagnosed as Atypical Rett Syndrome: Cautions in Interpretation of Genomic Test Results. *Semin Pediatr Neurol* **2018**, 26, 28–32. https://doi.org/10.1016/j.spen.2017.08.008.
(87) Chuan, Z.; Ruikun, C.; Qian, L.; Shiyue, M.; Shengju, H.; Yong, Y.; Haibo, L.; Neng, X.; Yong, Z.; Huiqin, X.; Weijia, W.; Ling, H.; Bingbo, Z.; Zhang, Q.; Yan, W.; Zongfu, C.; Xu, M. Genetic and Phenotype Analysis of a Chinese Cohort of Infants and Children With Epilepsy. *Front. Genet.* **2022**, 13. https://doi.org/10.3389/fgene.2022.869210.
(88) Caswell, R. C.; Gunning, A. C.; Owens, M. M.; Ellard, S.; Wright, C. F. Assessing the Clinical Utility of Protein Structural Analysis in Genomic Variant Classification: Experiences from a Diagnostic Laboratory. *Genome Med* **2022**, 14, 77. https://doi.org/10.1186/s13073-022-01082-2.
(89) Lasa-Aranzasti, A.; Larasati, Y. A.; Cardoso, J. da S.; Solis, G. P.; Koval, A.; Cazurro-Gutiérrez, A.; Ortigoza-Escobar, J. D.; Miranda, M. C.; Casa-Fages, B. D. la; Moreno-Galdó, A.; Tizzano, E. F.; Gómez-Andrés, D.; Verdura, E.; Katanaev, V. L.; Pérez-Dueñas, B.; Spain, S. G. of G. patients from. Clinical and Molecular Profiling in GNAO1 Permits Phenotype–Genotype Correlation. *Movement Disorders n/a* (n/a). https://doi.org/10.1002/mds.29881.
(90) Domínguez-Carral, J.; Ludlam, W. G.; Segarra, M. J.; Marti, M. F.; Balsells, S.; Muchart, J.; Petrović, D. Č.; Espinoza, I.; Ortigoza-Escobar, J. D.; Martemyanov, K. A.; Group, G.-S. Severity of GNAO1-Related Disorder Correlates with Changes in G-Protein Function. *Ann. Neurol.* **2023**, n/a (n/a), 987–1004. https://doi.org/10.1002/ana.26758.
(91) Feng, H.; Sjögren, B.; Karaj, B.; Shaw, V.; Gezer, A.; Neubig, R. R. Movement Disorder in GNAO1 Encephalopathy Associated with Gain-of-Function Mutations. *Neurology* **2017**, 89 (8), 762–770. https://doi.org/10.1212/wnl.0000000000004262.
(92) Knight, K. M.; Obarow, E. G.; Wei, W.; Mani, S.; Esteller, M. I.; Cui, M.; Ma, N.; Martin, S. A.; Brinson, E.; Hewitt, N.; Soden, G. M.; Logothetis, D. E.; Vaidehi, N.; Dohlman, H. G. Molecular Annotation of G Protein Variants in a Neurological Disorder. *Cell Reports* **2023**, 42 (12), 113462. https://doi.org/10.1016/j.celrep.2023.113462.
(93) Knight, K. M.; Krumm, B. E.; Kapolka, N. J.; Ludlam, W. G.; Cui, M.; Mani, S.; Prytkova, I.; Obarow, E. G.; Lefevre, T. J.; Wei, W.; Ma, N.; Huang, X.-P.; Fay, J. F.; Vaidehi, N.; Smrcka, A. V.; Slesinger, P. A.; Logothetis, D. E.; Martemyanov, K. A.; Roth, B. L.; Dohlman, H. G. A Neurodevelopmental Disorder Mutation Locks G Proteins in the Transitory Pre-Activated State. *Nat. Commun.* **2024**, 15 (1), 6643. https://doi.org/10.1038/s41467-024-50964-z.
(94) Solis, G. P.; Koval, A.; Valnohova, J.; Kazemzadeh, A.; Savitsky, M.; Katanaev, V. L. Neomorphic Gao Mutations Gain Interaction with Ric8 Proteins in GNAO1 Encephalopathies. *J. Clin. Investig.* **2024**, 134 (15), e172057. https://doi.org/10.1172/jci172057.
(95) Larasati, Y. A.; Solis, G. P.; Koval, A.; François-Heude, M.; Piarroux, J.; Roubertie, A.; Yang, R.; Zhang, Y.; Cao, D.; Korff, C. M.; Katanaev, V. L. Novel Mutation at Cys225 in GNAO1-Associated Developmental and Epileptic Encephalopathies: Clinical, Molecular, and Pharmacological Profiling of Case Studies. *MedComm (2020)* **2025**, 6 (5), e70196. https://doi.org/10.1002/mco2.70196.

(96) Larrivee, C. L.; Feng, H.; Quinn, J. A.; Shaw, V. S.; Leipprandt, J. R.; Demireva, E. Y.; Xie, H.; Neubig, R. R. Mice with GNAO1 R209H Movement Disorder Variant Display Hyperlocomotion Alleviated by Risperidone. *J. Pharmacol. Exp. Ther.* **2020**, 373 (1), 24–33. https://doi.org/10.1124/jpet.119.262733.
(97) Silachev, D.; Koval, A.; Savitsky, M.; Padmasola, G.; Quairiaux, C.; Thorel, F.; Katanaev, V. L. Mouse Models Characterize GNAO1 Encephalopathy as a Neurodevelopmental Disorder Leading to Motor Anomalies: From a Severe G203R to a Milder C215Y Mutation. *Acta Neuropathol Commun* **2022**, 10, 9. https://doi.org/10.1186/s40478-022-01312-z.

Table S2. Variant mechanism with citations.

| ***GNAO1* variant** | **Mechanism** | **Reference** |
| --- | --- | --- |
| G40R | Haplo | 90, 91, this paper |
| K278del | Haplo | This paper |
| Y231C | Haplo | 89,90, 92 |
| A227V | Haplo | 92 |
| S229R | Haplo | 92 |
| F275S | Haplo | 91 |
| L199P | Haplo | 90, 91, this paper |
| Y291N | Haplo | 89, 90,92 |
| G40W | Haplo | 92 |
| G40E | Haplo | 92 |
| D328G | Haplo | 92 |
| D273V | Haplo | 92 |
| D174G | Haplo | 91 |
| R179G | Haplo | 1 |
| N270D | Haplo | 92 |
| N270H | Haplo | 92 |
| G45R | Haplo | 92 |
| M244R | Haplo | 92 |
| F288S | Haplo | 92 |
| D273V | Haplo | 92 |
| Q52P | Haplo | 92 |
| Q52R | Haplo | 92 |
| D174N | Haplo | 92 |
| D273Y | Haplo | 92 |
| N270K | Haplo | 92 |
| K46E | Haplo | 92, 93, this paper |
| Q233P | Haplo | 94 |
| C225Y | Haplo | 95 |
| I279N | Haplo | 91 |
| G45E | Haplo | 92 |
| T48I | Haplo | 90 |
| G40E | Haplo | 92 |
| T182I | Haplo | 89,90 |
| R209C | Sequester G$\beta\gamma$ | 92, this paper |
| R209H | Sequester G$\beta\gamma$ | 92, 96, this paper |
| G203R | Sequester G$\beta\gamma$ | 92 |
| E237K | Sequester G$\beta\gamma$ | 92 |
| E246K | Sequester G$\beta\gamma$ | 92 |
| I344del | Sequester G$\beta\gamma$ | 89 |
| C215Y | Sequester G$\beta\gamma$ | 97 |
| A221D | Sequester G$\beta\gamma$ | 92 |
| R209L | Sequester G$\beta\gamma$ | 92 |
| D201V | Sequester G$\beta\gamma$ | 92 |
| G42R | Sequester G$\beta\gamma$ | 91 |
| G204R | Sequester G$\beta\gamma$ | 92 |
| R209G | Sequester G$\beta\gamma$ | 92 |
| E246Q | Sequester G$\beta\gamma$ | 92 |
| E246G | Sequester G$\beta\gamma$ | 92 |
| R209P | Sequester G$\beta\gamma$ | 90 |
| S207Y | Sequester G$\beta\gamma$ | 92 |
| L235P | Sequester G$\beta\gamma$ | 90 |
| S47G | Sequester G$\beta\gamma$ | 92 |
| S47N | Sequester G$\beta\gamma$ | 92 |
| K46R | Sequester G$\beta\gamma$ | 90 |
| G184R | Sequester G$\beta\gamma$ | 92 |
| L235P | Sequester G$\beta\gamma$ | 90 |

Table S3. Logistic regression of NEDIM status on mechanism and sex (full model with interaction).

| Term | $\beta$ (log-odds) | SE | OR | 95% CI | z | p |
| --- | --- | --- | --- | --- | --- | --- |
| Intercept (Sequester G$\beta\gamma$, F) | 1.338 | 0.28<br>1 | 3.8<br>1 | — | 4.77 | <.00<br>1 |
| Mechanism (Haplo vs. Sequester G$\beta\gamma$) | -2.948 | 0.5 | 0.0<br>5 | [0.02, 0.14] | -<br>5.89 | <.00<br>1 |
| Sex (M vs. F) | 0.22 | 0.42<br>4 | 1.2<br>5 | [0.54, 2.86] | 0.52 | 0.60<br>4 |
| Mechanism × Sex | 0.525 | 0.72<br>7 | 1.6<br>9 | [0.41, 7.03] | 0.72 | 0.47<br>1 |

N = 215. Reference categories: mechanism = Sequester G$\beta\gamma$, sex = F. Model fit: pseudo R<sup>2</sup> (McFadden) = 0.254, log-likelihood = -106.56 (null = -142.92), LR test p = 1.12e-15.

Table S4. Omnibus Type III tests — Figure 4 (membrane properties) outcomes

| Outcome | Scale | Term | F | df1 | df2 | P |
| --- | --- | --- | --- | --- | --- | --- |
| RMP | raw | genotype | 0.09 | 1 | 28.1 | 0.763 |
| RMP | raw | sex | 0.9 | 1 | 28.1 | 0.35 |
| RMP | raw | cell:type | 0.08 | 1 | 81.6 | 0.777 |
| RMP | raw | genotype:sex | 0.44 | 1 | 28.1 | 0.511 |
| RMP | raw | genotype:cell:type | 1.42 | 1 | 81.6 | 0.237 |
| RMP | raw | sex:cell:type | 0.9 | 1 | 81.6 | 0.347 |
| RMP | raw | genotype:sex:cell:type | 4.1 | 1 | 81.6 | 0.046 |
| Voltage deflection | raw | genotype | 40.49 | 1 | 28 | 6.93E-07 |
| Voltage deflection | raw | sex | 0 | 1 | 28 | 0.989 |
| Voltage deflection | raw | cell:type | 4.47 | 1 | 55 | 0.039 |
| Voltage deflection | raw | current | 495.15 | 1 | 83 | <0.0001 |
| Voltage deflection | raw | genotype:sex | 1.03 | 1 | 28 | 0.319 |
| Voltage deflection | raw | genotype:cell:type | 1.53 | 1 | 55 | 0.221 |
| Voltage deflection | raw | sex:cell:type | 1.08 | 1 | 55 | 0.303 |
| Voltage deflection | raw | genotype:current | 17.38 | 1 | 83 | 7.47E-05 |
| Voltage deflection | raw | sex:current | 0.07 | 1 | 83 | 0.798 |
| Voltage deflection | raw | cell:type:current | 1.26 | 1 | 83 | 0.265 |
| Voltage deflection | raw | genotype:sex:cell:type | 2.52 | 1 | 55 | 0.118 |
| Voltage deflection | raw | genotype:sex:current | 0.01 | 1 | 83 | 0.942 |
| Voltage deflection | raw | genotype:cell:type:current | 0.01 | 1 | 83 | 0.929 |
| Voltage deflection | raw | sex:cell:type:current | 0.01 | 1 | 83 | 0.928 |
| Voltage deflection | raw | genotype:sex:cell:type:current | 1.7 | 1 | 83 | 0.196 |
| Input resistance | log | genotype | 40.76 | 1 | 91 | 7.13E-09 |
| Input resistance | log | sex | 0.02 | 1 | 91 | 0.88 |
| Input resistance | log | cell:type | 5.47 | 1 | 91 | 0.022 |
| Input resistance | log | genotype:sex | 0.91 | 1 | 91 | 0.342 |
| Input resistance | log | genotype:cell:type | 1.95 | 1 | 91 | 0.166 |
| Input resistance | log | sex:cell:type | 0.24 | 1 | 91 | 0.624 |
| Input resistance | log | genotype:sex:cell:type | 2.46 | 1 | 91 | 0.121 |
| Membrane resistance | log | genotype | 34.86 | 1 | 30.3 | 1.75E-06 |
| Membrane resistance | log | sex | 5.6 | 1 | 30.3 | 0.025 |
| Membrane resistance | log | cell:type | 15.65 | 1 | 72.4 | 0.00018 |
| Membrane resistance | log | genotype:sex | 2.78 | 1 | 30.3 | 0.105 |
| Membrane resistance | log | genotype:cell:type | 15.15 | 1 | 72.4 | 0.00022 |
| Membrane resistance | log | sex:cell:type | 6.36 | 1 | 72.4 | 0.014 |
| Membrane resistance | log | genotype:sex:cell:type | 1.71 | 1 | 72.4 | 0.195 |
| Capacitance | log | genotype | 4.45 | 1 | 27.9 | 0.044 |
| Capacitance | log | sex | 3.75 | 1 | 27.9 | 0.063 |
| Capacitance | log | cell:type | 1.48 | 1 | 69.9 | 0.228 |
| Capacitance | log | genotype:sex | 1 | 1 | 27.9 | 0.327 |
| Capacitance | log | genotype:cell:type | 0.47 | 1 | 69.9 | 0.494 |
| Capacitance | log | sex:cell:type | 0.4 | 1 | 69.9 | 0.527 |
| Capacitance | log | genotype:sex:cell:type | 3.73 | 1 | 69.9 | 0.057 |

Linear mixed-effects models (mouse as random intercept; cell nested within mouse for repeated-current outcomes). "Scale" indicates raw units or log transformation. current terms apply only to outcomes with a repeated within-cell current-step factor.

Table S5. Pairwise genotype contrasts— Figure 4 (membrane properties) outcomes.

| Outcome | Scale | Sex | Cell type | Current (pA) | Estimate | t | df | P |
| --- | --- | --- | --- | --- | --- | --- | --- | --- |
| RMP | raw | F | D1 | — | -1.565 | -0.48 | 60.7 | 1 |
| RMP | raw | F | D2 | — | 0.252 | 0.08 | 59.3 | 1 |
| RMP | raw | M | D1 | — | 5.262 | 1.73 | 56.9 | 0.358 |
| RMP | raw | M | D2 | — | -1.74 | -0.53 | 72.4 | 1 |
| Voltage deflection | raw | F | D1 | -100 | -2.054 | -1.6 | 28 | 0.242 |
| Voltage deflection | raw | F | D1 | -50 | -1.045 | -0.95 | 28 | 0.35 |
| Voltage deflection | raw | F | D2 | -100 | -6.014 | -4.56 | 28 | 0.00019 |
| Voltage deflection | raw | F | D2 | -50 | -4.02 | -3.56 | 28 | 0.0014 |
| Voltage deflection | raw | M | D1 | -100 | -5.684 | -4.17 | 28 | 0.00053 |
| Voltage deflection | raw | M | D1 | -50 | -3.805 | -3.26 | 28 | 0.0029 |
| Voltage deflection | raw | M | D2 | -100 | -4.824 | -3.19 | 28 | 0.007 |
| Voltage deflection | raw | M | D2 | -50 | -3.805 | -2.94 | 28 | 0.007 |
| Input resistance | raw | F | D1 | — | 21.578 | 0.89 | 75.2 | 0.377 |
| Input resistance | raw | F | D2 | — | 82.57 | 3.37 | 80.9 | 0.0046 |
| Input resistance | raw | M | D1 | — | 75.457 | 3 | 78.7 | 0.011 |
| Input resistance | raw | M | D2 | — | 77.802 | 2.81 | 89.9 | 0.012 |
| Input resistance | log | F | D1 | — | 0.197 | 1.21 | 61.1 | 0.233 |
| Input resistance | log | F | D2 | — | 0.67 | 4.02 | 69.8 | 0.00058 |
| Input resistance | log | M | D1 | — | 0.6 | 3.52 | 72.2 | 0.0023 |
| Input resistance | log | M | D2 | — | 0.572 | 3 | 79.1 | 0.0072 |
| Membrane resistance | raw | F | D1 | — | 38.108 | 1.15 | 49.9 | 0.256 |
| Membrane resistance | raw | F | D2 | — | 81.777 | 2.47 | 48.3 | 0.051 |
| Membrane resistance | raw | M | D1 | — | 65.632 | 2.05 | 47.9 | 0.093 |
| Membrane resistance | raw | M | D2 | — | 211.721 | 6 | 64.6 | 3.94E-07 |
| Membrane resistance | log | F | D1 | — | 0.243 | 1.45 | 50.3 | 0.153 |
| Membrane resistance | log | F | D2 | — | 0.549 | 3.29 | 48.5 | 0.0056 |
| Membrane resistance | log | M | D1 | — | 0.399 | 2.47 | 48.2 | 0.034 |
| Membrane resistance | log | M | D2 | — | 1.015 | 5.65 | 66.6 | 1.44E-06 |
| Capacitance | raw | F | D1 | — | 19.943 | 1.9 | 47.9 | 0.253 |
| Capacitance | raw | F | D2 | — | 13.391 | 1.28 | 46.6 | 0.416 |
| Capacitance | raw | M | D1 | — | -4.064 | -0.4 | 46.1 | 0.691 |
| Capacitance | raw | M | D2 | — | 16.798 | 1.52 | 61.2 | 0.404 |
| Capacitance | log | F | D1 | — | 0.532 | 2.18 | 49.8 | 0.136 |
| Capacitance | log | F | D2 | — | 0.327 | 1.34 | 48.2 | 0.481 |
| Capacitance | log | M | D1 | — | -0.062 | -0.26 | 47.7 | 0.793 |
| Capacitance | log | M | D2 | — | 0.369 | 1.42 | 64.5 | 0.481 |

Estimated marginal means, Holm-corrected within each outcome/scale family. Estimate is on the model scale (raw units, or log-ratio for log-scale models).

Table S6. Omnibus Type III tests — Figure 5 (excitability) outcomes.

| Outcome | Scale | Term | F | df1 | df2 | P |
| --- | --- | --- | --- | --- | --- | --- |
| Rheobase | log | genotype | 23.92 | 1 | 103 | 3.72E-06 |
| Rheobase | log | sex | 1.34 | 1 | 103 | 0.249 |
| Rheobase | log | cell:type | 16.92 | 1 | 103 | 7.87E-05 |
| Rheobase | log | genotype:sex | 0.09 | 1 | 103 | 0.768 |
| Rheobase | log | genotype:cell:type | 3.54 | 1 | 103 | 0.063 |
| Rheobase | log | sex:cell:type | 0.15 | 1 | 103 | 0.704 |
| Rheobase | log | genotype:sex:cell:type | 2.4 | 1 | 103 | 0.125 |
| F-I curve (# APs) | raw | genotype | 49.69 | 1 | 30 | 7.77E-08 |
| F-I curve (# APs) | raw | sex | 1.01 | 1 | 30 | 0.323 |
| F-I curve (# APs) | raw | cell:type | 46.74 | 1 | 65 | 3.37E-09 |
| F-I curve (# APs) | raw | current | 241.03 | 7 | 665 | <0.0001 |
| F-I curve (# APs) | raw | genotype:sex | 0.05 | 1 | 30 | 0.819 |
| F-I curve (# APs) | raw | genotype:cell:type | 1.52 | 1 | 65 | 0.223 |
| F-I curve (# APs) | raw | sex:cell:type | 0 | 1 | 65 | 0.978 |
| F-I curve (# APs) | raw | genotype:current | 12.71 | 7 | 665 | 2.22E-15 |
| F-I curve (# APs) | raw | sex:current | 0.89 | 7 | 665 | 0.512 |
| F-I curve (# APs) | raw | cell:type:current | 8.22 | 7 | 665 | 1.26E-09 |
| F-I curve (# APs) | raw | genotype:sex:cell:type | 4.25 | 1 | 65 | 0.043 |
| F-I curve (# APs) | raw | genotype:sex:current | 0.1 | 7 | 665 | 0.998 |
| F-I curve (# APs) | raw | genotype:cell:type:current | 9.01 | 7 | 665 | 1.23E-10 |
| F-I curve (# APs) | raw | sex:cell:type:current | 0.88 | 7 | 665 | 0.523 |
| F-I curve (# APs) | raw | genotype:sex:cell:type:current | 1.21 | 7 | 665 | 0.296 |

Linear mixed-effects models (mouse as random intercept; cell nested within mouse for repeated-current outcomes). "Scale" indicates raw units or log transformation. current terms apply only to outcomes with a repeated within-cell current-step factor.

Table S7. Pairwise genotype contrasts — Figure 5 (excitability) outcomes.

| Outcome | Scale | Sex | Cell type | Current (pA) | Estimate | t | df | P |
| --- | --- | --- | --- | --- | --- | --- | --- | --- |
| Rheobase | raw | F | D1 | — | -18.529 | -0.65 | 55.3 | 0.516 |
| Rheobase | raw | F | D2 | — | -82.5 | -2.9 | 67.2 | 0.015 |
| Rheobase | raw | M | D1 | — | -84.856 | -3.23 | 63 | 0.008 |
| Rheobase | raw | M | D2 | — | -65.057 | -2.11 | 82.2 | 0.076 |
| Rheobase | log | F | D1 | — | -0.096 | -0.55 | 54 | 0.582 |
| Rheobase | log | F | D2 | — | -0.661 | -3.8 | 66.1 | 0.0013 |
| Rheobase | log | M | D1 | — | -0.4 | -2.48 | 62.1 | 0.047 |
| Rheobase | log | M | D2 | — | -0.454 | -2.4 | 81.2 | 0.047 |
| Rheobase | two-step (mouse-level) | F | D1 | — | — | 1.09 | — | 0.3 |
| Rheobase | two-step (mouse-level) | F | D2 | — | — | 3.33 | — | 0.0179 |
| Rheobase | two-step (mouse-level) | M | D1 | — | — | 2.99 | — | 0.0194 |
| Rheobase | two-step (mouse-level) | M | D2 | — | — | 3.63 | — | 0.0138 |
| F-I curve (# APs) | raw | F | D1 | 50 | 0 | 0 | 30 | 1 |
| F-I curve (# APs) | raw | F | D1 | 100 | -0.347 | -0.34 | 30 | 1 |
| F-I curve (# APs) | raw | F | D1 | 150 | -1.018 | -0.58 | 30 | 1 |
| F-I curve (# APs) | raw | F | D1 | 200 | 0.112 | 0.06 | 30 | 1 |
| F-I curve (# APs) | raw | F | D1 | 250 | 4.982 | 1.97 | 30 | 0.349 |
| F-I curve (# APs) | raw | F | D1 | 300 | 6.088 | 2.37 | 30 | 0.196 |
| F-I curve (# APs) | raw | F | D1 | 350 | 5.906 | 2.25 | 30 | 0.224 |
| F-I curve (# APs) | raw | F | D1 | 400 | 3.306 | 1.18 | 30 | 1 |
| F-I curve (# APs) | raw | F | D2 | 50 | 0.812 | 2.46 | 30 | 0.079 |
| F-I curve (# APs) | raw | F | D2 | 100 | 4.687 | 4.55 | 30 | 0.0005 |
| F-I curve (# APs) | raw | F | D2 | 150 | 9.487 | 5.38 | 30 | 6.31E-05 |
| F-I curve (# APs) | raw | F | D2 | 200 | 10.637 | 5.21 | 30 | 9.03E-05 |
| F-I curve (# APs) | raw | F | D2 | 250 | 6.775 | 2.65 | 30 | 0.064 |
| F-I curve (# APs) | raw | F | D2 | 300 | 2 | 0.77 | 30 | 0.448 |
| F-I curve (# APs) | raw | F | D2 | 350 | -3.7 | -1.39 | 30 | 0.348 |
| F-I curve (# APs) | raw | F | D2 | 400 | -6.388 | -2.25 | 30 | 0.097 |
| F-I curve (# APs) | raw | M | D1 | 50 | 0 | 0 | 30 | 1 |
| F-I curve (# APs) | raw | M | D1 | 100 | 0.846 | 0.89 | 30 | 0.765 |
| F-I curve (# APs) | raw | M | D1 | 150 | 2.462 | 1.51 | 30 | 0.426 |
| F-I curve (# APs) | raw | M | D1 | 200 | 5.322 | 2.81 | 30 | 0.051 |
| F-I curve (# APs) | raw | M | D1 | 250 | 7.091 | 2.99 | 30 | 0.043 |
| F-I curve (# APs) | raw | M | D1 | 300 | 7.231 | 3 | 30 | 0.043 |
| F-I curve (# APs) | raw | M | D1 | 350 | 6.245 | 2.54 | 30 | 0.083 |
| F-I curve (# APs) | raw | M | D1 | 400 | 4.413 | 1.68 | 30 | 0.417 |
| F-I curve (# APs) | raw | M | D2 | 50 | 0 | 0 | 30 | 1 |
| F-I curve (# APs) | raw | M | D2 | 100 | 2.545 | 2.28 | 30 | 0.18 |
| F-I curve (# APs) | raw | M | D2 | 150 | 6.255 | 3.27 | 30 | 0.021 |
| F-I curve (# APs) | raw | M | D2 | 200 | 6.827 | 3.08 | 30 | 0.03 |
| F-I curve (# APs) | raw | M | D2 | 250 | 4.955 | 1.79 | 30 | 0.336 |
| F-I curve (# APs) | raw | M | D2 | 300 | 1.409 | 0.5 | 30 | 1 |
| F-I curve (# APs) | raw | M | D2 | 350 | -3.118 | -1.08 | 30 | 0.862 |
| F-I curve (# APs) | raw | M | D2 | 400 | -6.2 | -2.01 | 30 | 0.267 |

Estimated marginal means, Holm-corrected within each outcome/scale family. Estimate is on the model scale (raw units, or log-ratio for log-scale models).

Table S8. F statistics for Figure 6.

| Panel | Sex | Drug | Source of Variation | F (DFn, DFd) | P value |
| --- | --- | --- | --- | --- | --- |
| C | Male | Gallein | Drug x Genotype | F (1, 15) = 13.12 | P=0.0025 |
|  |  |  | Drug | F (1, 15) = 42.69 | P<0.0001 |
|  |  |  | Genotype | F (1, 15) = 55.52 | P<0.0001 |
| D | Female | Gallein | Drug x Genotype | F (1, 18) = 33.71 | P<0.0001 |
|  |  |  | Drug | F (1, 18) = 78.57 | P<0.0001 |
|  |  |  | Genotype | F (1, 18) = 96.61 | P<0.0001 |
| E | Male | Caffeine | Drug x Genotype | F (1.449, 17.39) = 17.59 | P=0.0002 |
|  |  |  | Drug | F (1.449, 17.39) = 0.5093 | P=0.5516 |
|  |  |  | Genotype | F (1, 12) = 0.07109 | P=0.7943 |
| F | Female | Caffeine | Drug x Genotype | F (1.888, 30.21) = 11.56 | P=0.0002 |
|  |  |  | Drug | F (1.888, 30.21) = 6.982 | P=0.0037 |
|  |  |  | Genotype | F (1, 16) = 3.329 | P=0.0868 |

Table S9. Post hoc test p-values for Figure 6.

| Panel | Sex | Comparison type | Comparison | Adjusted p value |
| --- | --- | --- | --- | --- |
| C | Male | Between genotype | Vehicle: WT vs. R209H | <0.0001 |
| C | Male | Between genotype | Gallein: WT vs. R209H | 0.0001 |
| C | Male | Within genotype | WT: Vehicle vs. Gall ein | 0.0509 |
| C | Male | Within genotype | R209H: Vehicle vs. Gallein | <0.0001 |
| D | Female | Between genotype | Vehicle: WT vs. R209H | <0.0001 |
| D | Female | Between genotype | Gallein: WT vs. R209H | 0.1413 |
| D | Female | Within genotype | WT: Vehicle vs. Gall ein | 0.054 |
| D | Female | Within genotype | R209H: Vehicle vs. Gallein | <0.0001 |
| E | Male | Between genotype | Vehicle: WT vs. R209H | 0.0066 |
| E | Male | Between genotype | 25 mg/kg caffeine: WT vs. R209H | 0.2119 |
| E | Male | Between genotype | 50 mg/kg caffeine: WT vs. R209H | 0.2249 |
| E | Male | Within genotype | WT: Vehicle vs. 25 mg/kg caffeine | 0.0067 |
| E | Male | Within genotype | WT: Vehicle vs. 50 mg/kg caffeine | 0.0997 |
| E | Male | Within genotype | WT: 25 mg/kg caffeine vs. 50 mg/kg caffeine | 0.9339 |
| E | Male | Within genotype | R209H: Vehicle vs. 25 mg/kg caffeine | 0.0082 |
| E | Male | Within genotype | R209H: Vehicle vs. 50 mg/kg caffeine | 0.0285 |
| E | Male | Within genotype | R209H: 25 mg/kg caffeine vs. 50 mg/kg caffeine | 0.5268 |
| F | Female | Between genotype | Vehicle: WT vs. R209H | 0.0009 |
| F | Female | Between genotype | 25 mg/kg caffeine:<br>WT vs. R209H | 0.8741 |
| F | Female | Between genotype | 50 mg/kg caffeine:<br>WT vs. R209H | 0.2919 |
| F | Female | Within genotype | WT: Vehicle vs. 25<br>mg/kg caffeine | 0.0068 |
| F | Female | Within genotype | WT: Vehicle vs. 50<br>mg/kg caffeine | 0.3434 |
| F | Female | Within genotype | WT: 25 mg/kg<br>caffeine vs. 50 mg/kg<br>caffeine | 0.0043 |
| F | Female | Within genotype | R209H: Vehicle vs.<br>25 mg/kg caffeine | 0.2817 |
| F | Female | Within genotype | R209H: Vehicle vs.<br>50 mg/kg caffeine | 0.0049 |
| F | Female | Within genotype | R209H: 25 mg/kg<br>caffeine vs. 50 mg/kg<br>caffeine | 0.0539 |

Table S10. F statistics for Figure S5.

| Panel | Sex | Drug | Source of Variation | F (DFn, DFd) | P value |
| --- | --- | --- | --- | --- | --- |
| C | Male | Gallein | Drug x Genotype | F (1, 13) = 1.037 | P=0.3271 |
|  |  |  | Drug | F (1, 13) = 0.03827 | P=0.8479 |
|  |  |  | Genotype | F (1, 13) = 2.410 | P=0.1446 |
| D | Female | Gallein | Drug x Genotype | F (1, 12) = 8.645 | P=0.0124 |
|  |  |  | Drug | F (1, 12) = 10.39 | P=0.0073 |
|  |  |  | Genotype | F (1, 12) = 0.7238 | P=0.4115 |
| E | Male | KW-6002 and DPCPX | Drug x Genotype | F (1.538, 19.99) = 1.625 | P=0.2226 |
|  |  |  | Drug | F (1.538, 19.99) = 3.309 | P=0.0682 |
|  |  |  | Genotype | F (1, 13) = 5.888 | P=0.0305 |
| F | Female | KW-6002 and DPCPX | Drug x Genotype | F (1.461, 17.54) = 0.2655 | P=0.7007 |
| | | | Drug | $F(1.461, 17.54) = 3.354$ | $P=0.0708$ |
| | | | Genotype | $F(1, 12) = 13.86$ | $P=0.0029$ |

Table S11. Post hoc test p-values for Figure S5.

| Panel | Sex | Comparison type | Comparison | Adjusted p value |
| --- | --- | --- | --- | --- |
| C | Male | Between genotype | Vehicle: WT vs. R209H | 0.0768 |
| C | Male | Between genotype | Gallein: WT vs. R209H | 0.6014 |
| C | Male | Within genotype | WT: Vehicle vs. Gallein | 0.4209 |
| C | Male | Within genotype | R209H: Vehicle vs. Gallein | 0.5574 |
| D | Female | Between genotype | Vehicle: WT vs. R209H | 0.0405 |
| D | Female | Between genotype | Gallein: WT vs. R209H | 0.5059 |
| D | Female | Within genotype | WT: Vehicle vs. Gallein | 0.8169 |
| D | Female | Within genotype | R209H: Vehicle vs. Gallein | 0.0023 |
| E | Male | Between genotype | Vehicle: WT vs. R209H | 0.0368 |
| E | Male | Between genotype | KW-6002: WT vs. R209H | 0.0849 |
| E | Male | Between genotype | DPCPX: WT vs. R209H | 0.5301 |
| E | Male | Within genotype | WT: Vehicle vs KW-6002 | 0.2795 |
| E | Male | Within genotype | WT: Vehicle vs. DPCPX | 0.6115 |
| E | Male | Within genotype | WT: KW-6002 vs. DPCPX | 0.0703 |
| E | Male | Within genotype | R209H: Vehicle vs KW-6002 | 0.7325 |
| E | Male | Within genotype | R209H: Vehicle vs. DPCPX | 0.4241 |
| E | Male | Within genotype | R209H: KW-6002 vs. DPCPX | 0.0348 |
| F | Female | Between genotype | Vehicle: WT vs. R209H | 0.0489 |
| F | Female | Between genotype | KW-6002: WT vs. R209H | 0.0478 |
| F | Female | Between genotype | DPCPX: WT vs. R209H | 0.1869 |
| F | Female | Within genotype | WT: Vehicle vs KW-6002 | 0.1251 |
| F | Female | Within genotype | WT: Vehicle vs. DPCPX | 0.0589 |
| F | Female | Within genotype | WT: KW-6002 vs. DPCPX | 0.7861 |
| F | Female | Within genotype | R209H: Vehicle vs KW-6002 | 0.7719 |
| F | Female | Within genotype | R209H: Vehicle vs. DPCPX | 0.7287 |
| F | Female | Within genotype | R209H: KW-6002 vs. DPCPX | 0.8009 |
